# Probabilistic Cause-of-disease Assignment using Case-control Diagnostic Tests: A Latent Variable Regression Approach

**DOI:** 10.1101/672808

**Authors:** Zhenke Wu, Irena Chen

## Abstract

Optimal prevention and treatment strategies for a disease of multiple causes, such as pneumonia, must be informed by the population distribution of causes among cases, or cause-specific case fractions (CSCFs). CSCFs may further depend on additional explanatory variables. Existing methodological literature in disease etiology research does not fully address the regression problem, particularly under a case-control design. Based on multivariate binary non-gold-standard diagnostic data and additional covariate information, this paper proposes a novel and unified regression modeling framework for estimating covariate-dependent CSCF functions in case-control disease etiology studies. The model leverages critical control data for valid probabilistic cause assignment for cases.We derive an efficient Markov chain Monte Carlo algorithm for flexible posterior inference. We illustrate the inference of CSCF functions using extensive simulations and show that the proposed model produces less biased estimates and more valid inference of the overall CSCFs than analyses that omit covariates. A regression analysis of pediatric pneumonia data reveals the dependence of CSCFs upon season, age, HIV status and disease severity. The paper concludes with a brief discussion on model extensions that may further enhance the utility of the regression model in disease etiology research.

## 1 | INTRODUCTION

### 1.1 | Motivating Application

Pneumonia is a clinical condition associated with infection of the lung tissue, which can be caused by more than 30 different species of pathogens. In studies of pneumonia etiology, a cause is the subset of one or more pathogens infecting the lung. Knowledge about population-level cause-specific etiologic contributions can help prioritize prevention programs and design treatment algorithms. The Pneumonia Etiology Research for Child Health (PERCH) study is a seven-country case-control study of the etiology of severe and very severe pneumonia.^1^ The primary aim of the study is to estimate the etiologic contributions quantified by *cause-specific case fractions* (CSCFs), which may vary by individual-level factors such as age, enrollment date, disease severity, nutrition status and human immunodeficiency virus (HIV) status. We will refer to the covariate-dependent CSCFs as *CSCFfunctions*.

### 1.2 | Inferential Challenges: Imperfect Measurements and Additional Control Data

The fundamental challenge to estimating CSCFs is due to the unobservable true causes of disease among cases defined by clinical (non-microbiological) criteria. In PERCH study, tabulating case frequencies by cause is infeasible, because the lung-infecting pathogen(s) can rarely be directly observed due to potential clinical complications associated with invasive lung aspiration procedure.^2^ Alternatively, non-invasive real-time polymerase chain reaction (PCR) test was made on each case’s nasopharyngeal (NP) specimen (referred to as NPPCR), outputting presence or absence of a list of pathogens in the nasal cavity. The NP multivariate binary measurements are imprecise indicators for what infected the lung. In particular, detecting a pathogen in a case’s nasal cavity does not indicate it caused lung infection. To provide statistical control for false positive detections, PERCH study performed NPPCR tests on pneumonia-free controls.

Valid inference must take into account two salient characteristics: imperfect sensitivities and specificities. First, tests lacking sensitivity may miss the true causative pathogen(s) which if unadjusted may produce inferior estimates of the CSCF functions. Second, imperfect diagnostic specificities may result in false positive detection of pathogens that are not causes of pneumonia. Control data in PERCH study provide the requisite information for estimating false positive rates and must be integrated.

### 1.3 | Primary Aim: Regression Modeling with the Goal of Estimating CSCF Functions

Motivated by the case-control, non-gold-standard multivariate binary diagnostic test data from the PERCH study, statistical models and inferential algorithms have been successfully developed to estimate CSCFs without covariates.^3,4^ However, when individual-level explanatory variables are also available, the question of how to estimate CSCF functions remains. Data of similar structure have been collected by other large-scale disease etiology studies,^5,6^ raising acute needs for regression modeling.

The primary aim of our work is to leverage critical control diagnostic test data and covariate information when performing regression modeling of CSCFs among the cases, which is not studied before in the statistical literature. This is partly because such case-control studies of disease etiology using multiple diagnostic tests are completed only recently to provide new clinical and microbiological data to inform future prevention and treatment strategies. The models for no-covariate analysis are also developed recently, requiring extensions to regression settings to enable characterization of covariate-dependent disease etiology. See Section 7 for additional inferential challenges associated with small cell counts of cases and controls in covariate strata. To achieve this aim, we design a hierarchical Bayesian latent variable regression model for case-control multivariate binary responses and derive an efficient posterior inference algorithm. In addition, through extensive simulation studies, we demonstrate that our regression model produces more valid inference than competing case-control models that cannot use covariates.

The rest of the paper is organized as follows. Section 2 describes the data structure and introduces notations. Section 3 contextualizes our work by reviewing related literature. Section 4 reviews existing models without covariates. Section 5 formulates the proposed regression model, specifies priors, and derives the posterior sampling algorithm. We demonstrate the proposed method via extensive simulations in Section 6 and an application to PERCH data in Section 7. The paper concludes with a discussion on future research directions.

## 2 | DATA STRUCTURE AND NOTATIONS

Let *Y_i_* = 1 indicate a case subject with the clinically-defined disease and *Y_i_* = 0 indicate a control subject without disease. Let ***M**_i_* = (***M***_*i*1_,…***M***_*iJ*_)^T^ ∈ {0,1}^*J*^ represent the multivariate binary case-control, non-gold-standard diagnostic test results from subject *i*. Let 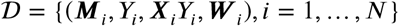 represent data, where ***X**_i_* = (*X*_*i*1_,…, *X_ip_*)^⊤^ are the *p* primary covariates in CSCF functions and hence must be available for cases, and ***W**_i_* = (*W*_*i*1_,…, *W_iq_*)^⊤^ are *q* covariates that are available in the cases and the controls. ***X**_i_* and ***W**_i_* may be identical, overlapping or completely different. ***X**_i_Y_i_ = **X**_i_* for a case *Y_i_* = 1; ***X**_i_Y_i_* is a vector of zeros for a control subject. For notational convenience, we have ordered the continuous variables, if any, in ***X**_i_* and ***W**_i_* as the first *p*_1_ and *q*_1_ elements, respectively. In this paper, we focus on pre-specified ***X**_i_* and ***W**_i_* and discuss the important problem of variable selection in Section 8.

## 3 | RELATED LITERATURE

The scientific problem of estimating CSCFs can be naturally formulated as estimating the mixing weights in a finite-mixture model where the mixture components represent distinct data generating mechanisms under different causes of diseases. In this paper, “cause” does not represent an intervention as in classical causal inference literature. Rather, it describes a specific mechanism leading up to the disease and follows the use in demography and disease etiology.^7,3,8^

### Case-only Methods

A closely related application in demography is to use verbal autopsy (VA) surveys to estimate the cause-specific mortality fractions (CSMFs) in regions without vital registry. Early methods rely on gold-standard cause-of-death information^7,8^ proposed an unsupervised, informative Bayes implementation of a latent class model (“LCMs”),^9^ where the latent classes represent unobserved causes of death and the survey responses are mutually independent given each latent class. Moran et al.^10^ let covariates influence the conditional distribution of the survey responses given a cause via hierarchical factor regression models. However, these methods do not account for epidemiological factors and individual characteristics that may influence the CSMFs. Datta et al.^11^ recognized the variation of CSMFs by covariates and studied transfer learning from a source population to a target population with a few deaths with observed causes. Finally, VA data are by definition case-only, hence are not applicable in our case-control setting.

### Case-control Latent Variable Methods

Methods that use *case-control* data to estimate CSCFs remain sparse. We review the only existing methods. Wu et al.^3^ introduced a *partially-latent class model* (pLCM) as an extension to classical LCMs. In particular, the pLCM is a *semi-supervised* method: it assumes with probabilities ***π*** = (*π*_1_,…, *π_L_*)^⊤^, or CSCFs, a case observation is drawn from a mixture of *L* components each representing a cause of disease, or “disease class”. Controls have no infection in the lung hence are drawn from an observed class. Each causative pathogen is assumed to be observed with a higher probability (sensitivity, or true positive rate, TPR) in case class *ℓ* than among the controls. A non-causative pathogen is observed with the same probability as in the controls (1 - specificity, or false positive rate, FPR). Under the pLCM, a higher observed marginal positive rate for a pathogen among cases than controls indicates its etiologic importance. Bayes rule is used to estimate ***π*** and other parameters via simple Markov chain Monte Carlo (MCMC) algorithms. The latent variable formulation has the unique practical advantage of integrating information from multiple data sources, including extra case-only data and multiple case-control measures, to estimate individual-level probabilities. Like classical LCMs, the pLCM assumes “local independence” (LI) which means the measurements (*M*_*i*1_,…, *M_iJ_*) are mutually independent given the disease class membership. Deviations from LI, or “local dependence” (LD) are testable using the control data, which is modeled by *nested* pLCM (npLCM)^4^ to reduce the bias in CSCF estimation. The importance of modeling LD to reduce the biases in estimating the mixing weights and response probabilities has also been extensively studied before but in the context of classical LCMs.^12,13^ Finally, the npLCM is partially-identified,^4^ necessitating informative priors for a subset of parameters (TPRs), which in PERCH study are obtained from external vaccine probe studies.

## 4 | PRELIMINARIES: EXISTING MODELS WITHOUT COVARIATES

### 4.1 | Additional Notations for Unobserved Causes-of-disease: “Latent States”

We first introduce notations for the unobserved causes of disease for the case subjects. Suppose a total of *J* “agents” or “items” are measured by the diagnostic tests. Let a binary variable *ι_ij_* indicate whether or not the *j*-th agent caused case *i*’s disease, that is, *ι_ij_* = 1 if *j*-th agent caused the disease, *ι_ij_* = 0 otherwise. We also allow more than one agent to cause the disease. We therefore have ***ι**_i_* = (*ι*_*i*1_,…, *ι_iJ_*)^⊤^ ∈ {0,1}^*J*^ which is a vector of multiple binary indicators that represent the causes for subject *i*. We will also refer to ***ι**_i_* as “latent states” for case subject *i*. Note that we allow the all-zero latent states ***ι**_i_* = 0_*J*×1_. This is to represent a case subject with a “Not Specified” (NoS) cause which, in PERCH study, represents the subgroup of cases whose diseases are caused by agents not specified as molecular targets in the diagnostic PCR tests. Cases can thus be classfied by distinct multivariate binary patterns of ***ι**_i_*. We will refer to cases having the same pattern of ***ι**_i_* as belonging to the same disease class.

In this paper, we assume that there are *L* classes of *pre-specified* latent state patterns (non-zero or all-zero) among the case subjects. Let a set 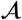 comprise the pre-specified *L* distinct multivariate binary patterns so that 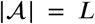, where 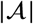 is the cardinality of 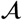. We discuss the important problem of unknown subsets in Section 8. We then introduce disease class indicators by arbitrarily labeling elements in 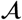 from 1 to *L*. We can now use *I_i_* that takes its value from {1,…, *L*} to indicate case subject *i*’s class. We also let 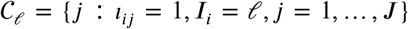 represent the subset of causative agents for disease class *ℓ*; for NoS class, we have 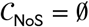.

For a control subject *i′* without disease, we use *I_i′_*, = 0 to indicate ***ι**_i′_*, = **0**_*J*×1_ which means no measured pathogen is a cause. For a case or a control subject, the value of *I_i_* thus corresponds to a particular latent state pattern; we denote this correspondence by ***ι**_i_ = **ι**_i_*(*I_i_*) which will be handy for specifying the models.

To illustrate the scientific meaning of the notations, consider a hypothetical list of *J* = 5 species of pathogens that are targeted by the NPPCR tests and possible disease-causing agents in the context of PERCH study. First, under an assumption of single-pathogen causes and no NoS class, we have *L = J* = 5 disease classes with distinct patterns of 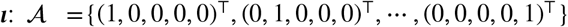. We can label the five disease classes by 1,…, *L* = 5, so that, for example, *I_i_* = 2 corresponds to ***ι**_i_* = (0,1,0,0,0)^⊤^ and 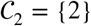, *I_i_* = 5 corresponds to ***ι**_i_* = (0,0,0,0,1)^⊤^ and 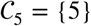. Second, under a less restrictive assumption of single- or double-pathogen causes (still no NoS class), we have 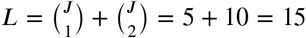 disease classes. For example, cases with the first and third pathogen infecting the lung are represented by ***ι**_i_* = (1,0,1,0,0)^⊤^ which has the subset of causes 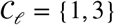 where *I_i_ = ℓ* is an arbitrary integer label of the disease class.

We first briefly review existing models without covariates, because they are the basis for our proposed regression model. We will describe the likelihood functions via generative processes, first for control data and then for case data. We then explain the rationales along with each step to provide scientific motivations for the model structures.

### 4.2 | PLCM: No covariate, conditional independence

We first review partially-latent class models (pLCM)^3^, which assumes that a subset of subjects have observed states. In PERCH, this means assuming the control subjects have no pathogen infecting the lungs. Although a control subject has no disease, positive responses among ***M**_i_* may result from imperfect specificities of the diagnostic tests. In particular, the data generating process for control data is from *J* independent Bernoulli trials with distinct success probabilities:

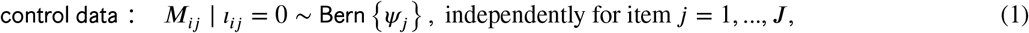

where the parameters ***ψ*** = (*ψ*_1_,…, *ψ_J_*)^⊤^ represent the positive response probabilities absent disease, referred to as “false positive rates” (FPRs), or 1-specificity. The data generating process for cases is an *L*-component finite mixture model. In the following, Step (2) generates a disease class indicator for a case subject that takes value from {1,…, *L*}. The lung-infecting pathogens are represented by ***ι**_i_*, which is found by the correspondence between the disease class indicator *I_i_* and ***ι**_i_*. Given ***ι**_i_*, Step (3) generates measurements of ***ι**_i_* resulting in error-prone multiple responses {*M_i_*_1_*,…, M_iJ_*} with positive response probabilities 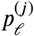 that takes the value of *θj* or *ψ*. according as whether the *j*-th agent caused the disease. We have

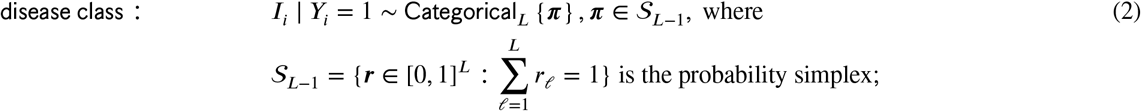

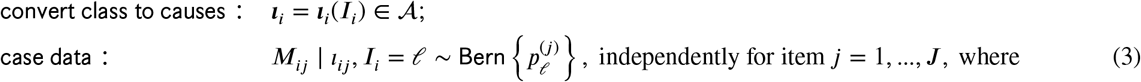

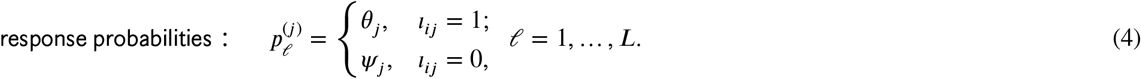

We term the parameter *θ_j_* as “true positive rate” (TPR), or sensitivity. It is assumed to be larger than the FPR *ψ_j_* (*θ_j_ > ψ_j_*). For the case model, the pLCM makes a key “non-interference” assumption in (4) that disease-causing pathogen(s) are more frequently detected among cases than controls and the non-causative pathogens are observed with the same rates among cases as in controls^4^. Under the single-pathogen-cause assumption, pLCM uses *J* TPRs ***θ*** = (*θ*_1_,…, *θ_J_*)^⊤^ and *J* FPRs ***ψ*** = (*ψ*_1_,…, *ψ_J_*)^⊤^. Note that (1) and (3) assume conditional independence of the measurements given the lung status *I_i_*; this is referred to as local independence (LI).

### 4.3 | Nested PLCM: No covariate, conditional dependence

The proposed regression model in this paper will build upon an extension to pLCM, referred to as “nested pLCM”.4We therefore provide a review of the npLCM model structure and rationale. The npLCM is designed to reduce potential estimation bias in ***π*** under large degrees of deviations from LI assumed in the original pLCM. It achieves the aim by characterizing residual correlations among *J* binary measurements ***M**_i_* even after conditioning on *I_i_*, e.g., the controls (*I_i_* = 0) or a case class (*I_i_ = ℓ, ℓ* ≠ 0).^4^ The extension is motivated by the flexibility of the classical LCM formulation^9^ to approximate any joint distribution of multivariate discrete responses.^14^

Compared to the pLCM, the data generating process for control data now assumes an additional step of generating “subclasses”. Scientifically, the subclasses in the controls (and cases in the following) can represent subgroups of children with different levels of immunity with positive rates varying across subclasses. Importantly, the conditional dependence of [***M**_i_* | *I_i_*] arises after the subclass indicators are integrated out. More specifically, we assume

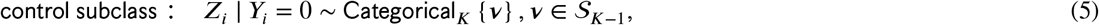

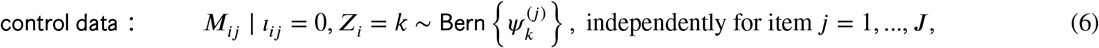

where ***ν*** = (*ν*_1_,…, *ν_K_*)^⊤^ is the vector of subclass probabilities and lies in a probability simplex. The original pLCM results if *K* = 1. Let 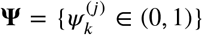 be a *J × K* matrix comprising FPRs, which are necessary for modeling the imperfect binary measurements among the controls. Let ***ψ***^(*J*)^ and ***ψ**_k_* represent the *j*-th row and *k*-th column. The data generating process for cases is as follows, with an additional Step (8) for drawing a subclass indicator *Z_i_* for each case subject:

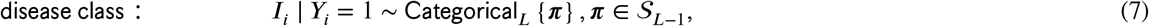

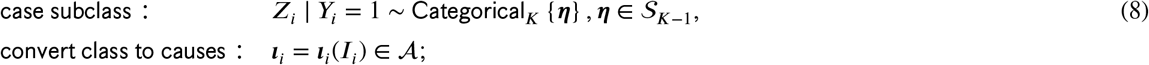

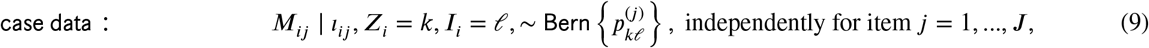

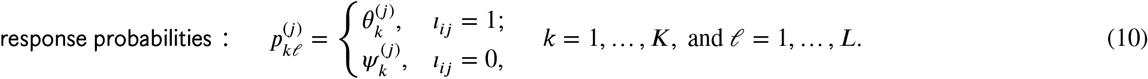

At Step (8), the npLCM introduces *K* unobserved subclasses with weights ***η*** = (*η*_1_,…, *η_K_*)^⊤^. The weights are shared across *L* disease classes. Let 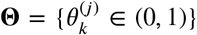 be a *J × K* matrix where 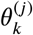 represents the positive response probability in subclass *k* if item *J* is causative in a disease class. We also refer to 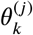 as TPR or sensitivity as in pLCM. Let ***θ***^(*J*)^ and ***θ**_k_* represent the *j*-th row and *k*-th column. In Step (10), 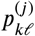 represents the positive response probability of *M_ij_* in subclass *k* of disease class *ℓ*, which equals the TPR 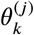 for a causative pathogen and the FPR 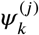 otherwise. We collect all the positive response probabilities for subclass *k* in disease class *ℓ* into 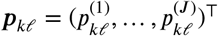.

#### 4.3.1 | Subclasses: Nuisance Parameters for Characterizing Residual Correlations Given *I_t_*

Different from what are of primary interest (the disease class indicators *I_i_* and its population distribution CSCFs ***π***), the latent subclasses *Z_i_* nested in each class *ℓ* = 0,1,…, *L* are *nuisance* parameters introduced to approximate complex multivariate dependence among discrete data. Importantly, the conditional dependence of [***M**_i_* | ***I**_i_*] arises after the subclass indicators are integrated out with respect to subclass weights (***ν*** or ***η***). In particular, Wu et al.^4^ used truncated stick-breaking priors for the subclass weights to encourage few subclasses and to side-step the choice of the true number of subclasses. However, nuisance parameter does not mean it is not important for improving the inference about the primary substantive parameters - the CSCFs. To the contrary, Wu et al.^4^ showed via asymptotic bias calculations and simulations that the use of subclasses reduces the estimation bias of CSCFs in the context without covariates and improves the coverage of the credible intervals.

Scientifically, the subclasses in the cases and controls can represent subgroups of children with different levels of immunity with positive rates ***θ**_k_* and ***ψ**_k_* varying across subclasses.

##### Remark 1.

Similar to the pLCM, the FPRs **Ψ** in the npLCM are shared between the controls and the case classes for noncausative agents (via 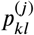). Different from the pLCM, the subclass mixing weights may differ between the cases (***η***) and the controls (***ν***). The special case of *η_k_* = *ν_k_, k* = 1,…, *K*, means the covariation patterns among the non-causative pathogens in a disease class is not different from the controls. However, relative to the controls, cases may have different strength and direction of observed pairwise measurement dependence in each disease class. By allowing the subclass weights to differ between the cases and the controls, npLCM is more flexible than pLCM in referencing cases’ measurements against the controls.

## 5 | PROPOSED REGRESSION EXTENSION

### 5.1 | Overview

We now describe the likelihood of the proposed regression model by a generative process building on npLCM. Only Steps (11), (12) and (13) below are results of the proposed extension; other parts of npLCM remains the same.

Data from controls provide requisite information about the specificities (1-FPRs) and covariations that may depend on covariates, which must be modeled for valid probabilistic cause assignment. The proposed model assumes the control subclass weights are covariate-dependent:

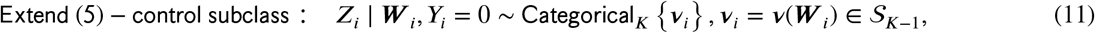

where, as in npLCM^4^, the subclass indicators *Z_i_*’s are nuisance quantities for inducing dependence among the multivariate binary responses ***M**_i_*, but now given covariates. ***ν**_i_* = (*ν_i_*_1_,…, *ν_iK_*) is the vector of control subclass probabilities that now may depend on ***W**_i_*. Scientifically, we are not interested in how the subclass probabilities are associated with covariates. We introduce ***ν***(***W***) here because, upon integrating over the distribution of *Z_i_* in (11), it helps define a flexible conditional distribution of ***M**_i_* given covariates ***W**_i_*.

FPR subclass profile *k*, 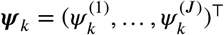, receives a weight of *ν_k_*(***W**_i_*) for a control subject *i* with covariates ***W**_i_*. After integrating out *Z_i_*, we obtain *marginal* control FPRs: 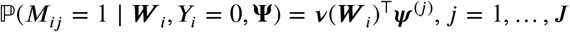, which depend on ***W**_i_*. For example, this enables us to use control data to estimate seasonal marginal FPRs of pathogen A. During seasons with frequent asymptomatic carriage among the controls, we may estimate the FPRs to be high. This means that presence of pathogen A in a case’s nasal cavity (the body site where the diagnostic tests in PERCH study were done) during these seasons does not necessarily indicate etiologic importance.

In Section 5.4.1, we propose a novel prior for the probability simplex regression ***ν***(***W**_i_*) to encourage fewer effective subclasses and side-step the choice of *K* by setting it to a large number that is appropriate for specific applications.

A reviewer raised the question of why formulating the model with subclass weights but not the TPRs and FPRs depending on covariates ***W**_i_*. Our modeling choice is primarily driven by observed conditional dependence among the multivariate binary measurements even after conditional on the covariates, which in the context of PERCH study means that diagnostic measurements of pathogens still exhibit mutual inhibition or stimulation even after stratifying on individual-level covariates, e.g, age. If we directly regressed TPRs and FPRs on covariates without introducing subclasses, a more restrictive assumption of independent measurements given covariates in each disease class would result. See Appendix A1 in the Supplementary Materials for further remarks on the control model assumption and the use of subclasses in this paper. In addition, the biological motivation for introducing subclasses is to characterize pathogen mutual inhibitions or stimulations by distinct the response profiles in the subclasses. The model structure based on subclasses is also amenable to posterior computations via data augmentation.

For cases, we follow the case model for npLCM, but extend in two aspects: let CSCFs depend on covariates ***X**_i_* and let case subclass weight depend on covariates ***W**_i_*. That is,

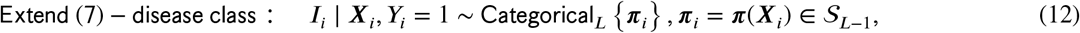

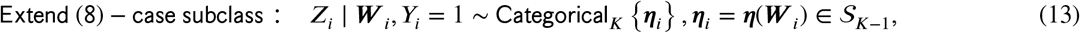

where ***π***(***X**_i_*) =(*π*_1_(***X**_i_*),…, *π_L_*(***X**_i_*))^⊤^ are CSCF functions evaluated at ***X**_i_*, and ***η***(***W**_i_*) = (*η*_*i*1_(***W**_i_*),…, *η_iK_*(***W**_i_*))^⊤^ is the vector of case subclass probabilities evaluated at ***W**_i_*. Both ***π**_i_* and ***η**_i_* are quantities from probability simplexes.

In this paper, we have adopted the approach of specifying an upper bound of working number of subclasses, and using sparsity priors to encourage towards a few subclasses for fitting the data. In Simulation I in Section 6, we illustrate a scenario where in truth two subclasses are present. During the model fitting, agnostic to the truth, we specify an upper bound of 7. Figure S1 in the Supplementary Materials shows that the model can learn from the data and figure out that two subclasses are actually needed. If the working number of subclasses *K* is too small, the model may not be flexible enough to characterize the dependence among the measurements. We therefore recommend starting with a number, e.g., 5 – 10, and then if computational resource is ample, trying larger *K*s. We usually observe that results remained similar for larger *K*. We can then report the final result based on a small *K* that gives stable results.

### 5.2 | Targets of Inference

In this paper, CSCF functions ***π***(***X***) is the primary target of inference. In addition, let 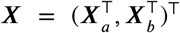, where ***X**_a_* and ***X**_b_* are *pre-specified* partition of ***X***. The partition is a generic notation, and is useful to represent, e.g., ***X**_b_* = {`I(age > 1), I(HIV positive), I(very severe pneumonia)`} and *X_a_* can be enrollment date. It is also of policy interest to infer the overall CSCFs given a specific age-HIV-severity stratum, while integrating over the study period.^1^ We define the *overall* CSCFs in a stratum ***X**_b_* = ***x**_b_* by integrating CSCF functions over the subset of covariates ***X**_a_* while holding ***X**_b_* fixed at specified covariate values ***x**_b_*:

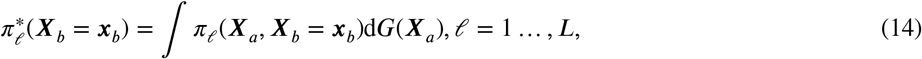

where *G* is a probability or empirical distribution of [***X**_a_ | X_b_*]. Here [*A | B*] represents the conditional distribution of random quantity *A* given another random quantity *B*. When *B* = Ø, it represents the distribution of *A*. We simply write 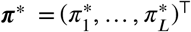 when ***X**_b_* = Ø. Following the definition, ***π***\* {`I(age > 1) =0, I(HIV positive) = 0, I(very severe pneumonia) = 0`} characterizes the overall CSCFs in the specific age-HIV-severity stratum. Inferences made about ***π***(***X***) and ***π***\*(***X**_b_*) are demonstrated in Section 7.

### 5.3 | Detailed Regression Specifications

We first specify the functional forms of ***π***(·), ***v***(·), ***η***(·), based on which we complete the joint distribution specification by describing the priors in Section 5.4.

#### 5.3.1 | Primary Component: CSCF Regression Model

We assume the CSCFs depend on ***X_i_*** via a classical multinomial logistic regression model:

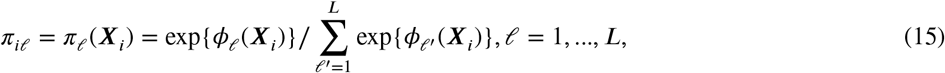

where *φ_ℓ_*(***X**_i_*) − *φ_L_*(***X**_i_*) is the log odds of case *i* in disease class *ℓ* relative to *L*: log *π_iℓ_/π_iL_*. The choice of not fixing a baseline category with zero regression coefficients is based on convenient prior specification so that all categories are treated symmetrically. Although the actual values of regression coefficients are not identifiable, contrasts between parameters in distinct categories remain identifiable, which are then summarized in the posterior inference. We further assume additivity in a partially linear model:

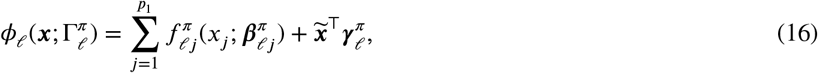

where 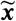 is the subvector of the predictors ***x*** that enters the model for all disease classes as linear predictors which may include an intercept, and 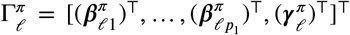 is the vector of the regression coefficients for disease class *ℓ*. Section 8 discusses non-additive extensions. For covariates such as enrollment date that serve as proxy for factors driven by seasonality, non-linear functional dependence is expected. In Section 5.4.2, we approximate unknown functions of a standardized continuous variable such as 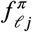 via basis expansions and along with a prior on the basis coefficients to encourage smoothness.

Integrating over *L* unobserved disease classes and *K* subclasses in (13-12), we obtain the likelihood for cases:

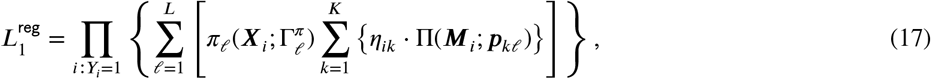

where 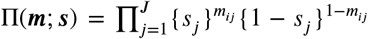 is the probability of observing *J* independent Bernoulli-distributed random variables with success probabilities ***s*** = (*s*_1_,…, *s_J_*)^⊤^∈ [0,1]^*J*^. In the following, we parameterize *η_ik_* by a logistic stick-breaking regression technique, which we first introduce in the control model.

#### 5.3.2 | Nuisance Components: Covariate-dependent Reference Distribution

The control data serve as reference against which diagnostic test results obtained from the cases are compared when estimating cause-specific probabilities. The control model is a mixture model with covariate-dependent mixing weights ***ν**_i_ = **ν***(***W**_i_*).

##### Control subclass weight regression

We specify *ν_ik_* by logistic stick-breaking parameterization:

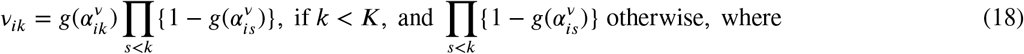

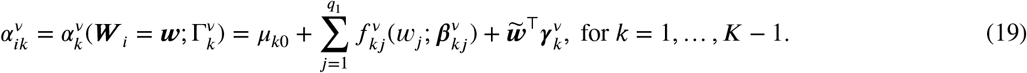

Let 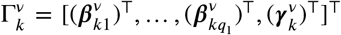 be the regression coefficients in the *k*-th subclass, and 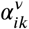 is subject *i*’s linear predictor at stick-breaking step *k* = 1,…, *K* − 1; *g*(·): 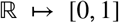 is a link function. In this paper, we use the logistic function *g*(*α*) = 1/{1 + exp(−*α*)} which is consistent with (15) so that the priors of the coefficients 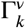 and 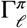 can be similar. In addition, the parameterization is amenable to simple and efficient block posterior sampling via Pólya-Gamma augmentation^15^. Generalization to other link functions such as the probit function is straightforward.^16^ Using the stick-breaking analogy, we begin with a unit-length stick: for a total of *K* − 1 stick-breaking events, we break a fraction 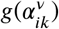 from the remaining stick at step *k*, resulting in segment *k* of length *ν_ik_*; this procedure is repeated for *k* = 1,…, *K* − 1.

It is readily derived that the likelihood for controls is: 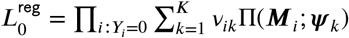.

##### Case subclass weight regression

The case subclass weight curve ***η_k_***(***W***) is also specified via a logistic stick-breaking regression as in the controls but with different linear predictors 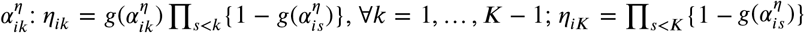. Given **Θ** and **Ψ**, ***η**_k_*(***W***) fully determines the joint distribution [***M** | **W, I*** = ℓ ≠ 0, **Θ, Ψ**]. We do not assume *η_k_*(***w***) = *ν_k_*(***w***), ∀***w***. Consequently, relative to the controls, the individuals in disease class *ℓ* may have different strength and direction of observed dependence between the causative 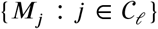 and non-causative 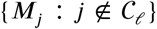 pathogens, or between the non-causative pathogens. Let the *k*-th linear predictor

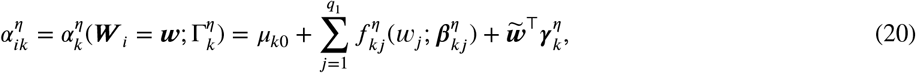

where 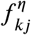 and 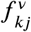 (from the control model) share the basis functions but the regression coefficients 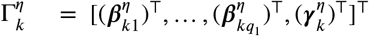 differ from the control counterpart 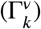. In addition, we have used the same intercepts {*μ*_*k*0_} in (19)to ensure only important subclasses in the controls are used in the cases. For example, absent covariates ***W***, a large and positive *μ*_*k*0_ effectively halts the stick-breaking procedure at step *k* for the controls. This is because the *k*-th stick-breaking will take almost the entire remaining stick, resulting in *ν*_*k*+1_ that is approximately zero. Applying the same intercept *μ*_*k*0_ to the cases makes *η*_*k*+1_ ≈ 0.

The joint likelihood for the proposed model is 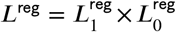. This completes the likelihood specification for the proposed model.

###### Remark 2.

Sometimes ignoring ***X**_i_* and ***W**_i_* does not invalidate inference of the overall CSCFs ***π***\*. For example, assuming covariate-independence: ∀*k, η_k_*(·) ≡ *η_k_*, 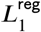 and 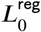 integrate to 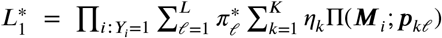, and 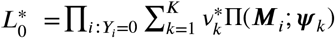, where 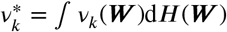 and *H* is probability or empirical distribution of ***W***. The integrated likelihood 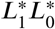 is now an npLCM likelihood without covariates. In addition, it can be readily shown that no-covariate inference for ***π***\* is also valid when ***X*** and ***W*** do not share common elements.

### 5.4 | Prior Distribution

The number of parameters in 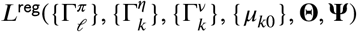 is 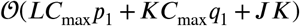 where *C*_max_ is the maximum number of basis functions in 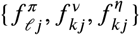. It easily exceeds the number of observed distinct binary measurement patterns. To overcome potential overfitting and increase model interpretability, we *a priori* encourage the following two features: (a) few non-trivial subclasses uniformly over ***W**_i_* values, and (b) constant subclass weights over ***W**_i_* values *η_k_*(·) = *η_k_* and *ν_k_*(·) = *ν_k_*.

#### 5.4.1 | Priors: Encourage Few Subclasses

Based on the parameterization of ***ν***(***W***) in (18) and (19), we propose a novel prior for ***ν***(***W***) over a probability simplex 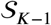. Because the linear predictor 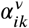 depends on *μ*_*k*0_ and other regression coefficients, priors on *μ*_*k*0_ and the coefficients induce a prior distribution for 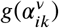 and thus a prior for *ν*(***W***) according to (18). For example, consider 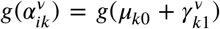, for subclass denoted as *k = A, B*. The prior for *μ*_*k*0_ and a Gaussian prior for 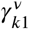 will induce a prior for 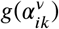 and thus for (*ν_A_, ν_B_, 1−ν_A_−ν_B_*)^⊤^.

Specifically, our basic idea is to have one of 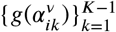 in (18) close to one *a posteriori* by making the posterior mean of one of 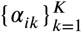 large. We accomplish this by specifying a novel additive prior on the intercept in (19):

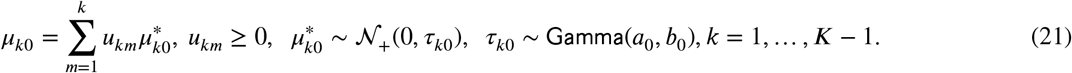

where {*u_km_, m* = 1,…, *k*, and *k* = 1,…, *K* − 1} is a pre-specified triangular array. In this paper, we use *u_km_* = 1, *m* = 1,…, *k*. Other choices, such as *u_km_* = **1**{*k = m*} or *u_km_* = 1/*k*, may be useful in other settings. Here, 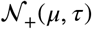 represents a Gaussian distribution with mean *μ*, precision *τ* truncated to the positive half. We set shape *a*_0_ = *ν*_0_/2, and rate 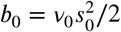. Marginalized over *τ*_*k*0_, 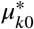 has a truncated scaled-*t* distribution with degree of freedom *ν*_0_ and scale *s*_0_, which peaks at zero and has a heavy right tail. A large positive value from the heavy-tailed prior for 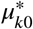 at stick-breaking step *k* produces a large *μ*_*k*0_ and takes away nearly the entire stick segment currently left. We will detail the Gaussian priors for other regression coefficients in the next two subsections.

For a fixed *ν*_0_, the scale parameter *s*_0_ modulates the tendency for the prior density of ***ν***(***W***) to concentrate towards a few vertices in a probability simplex (see Figure 2). Under the prior (21), it makes higher-order subclasses *a priori* increasingly unlikely to receive substantial weights. As a result, once applied to the control model (11), the prior encourages using a small number of subclasses to approximate the observed 2^*J*^ covariate-dependent probability contingency table for the control data {***M**_i_, **W**_i_*: *Y_i_* = 0} in finite samples.

**FIGURE 1.**
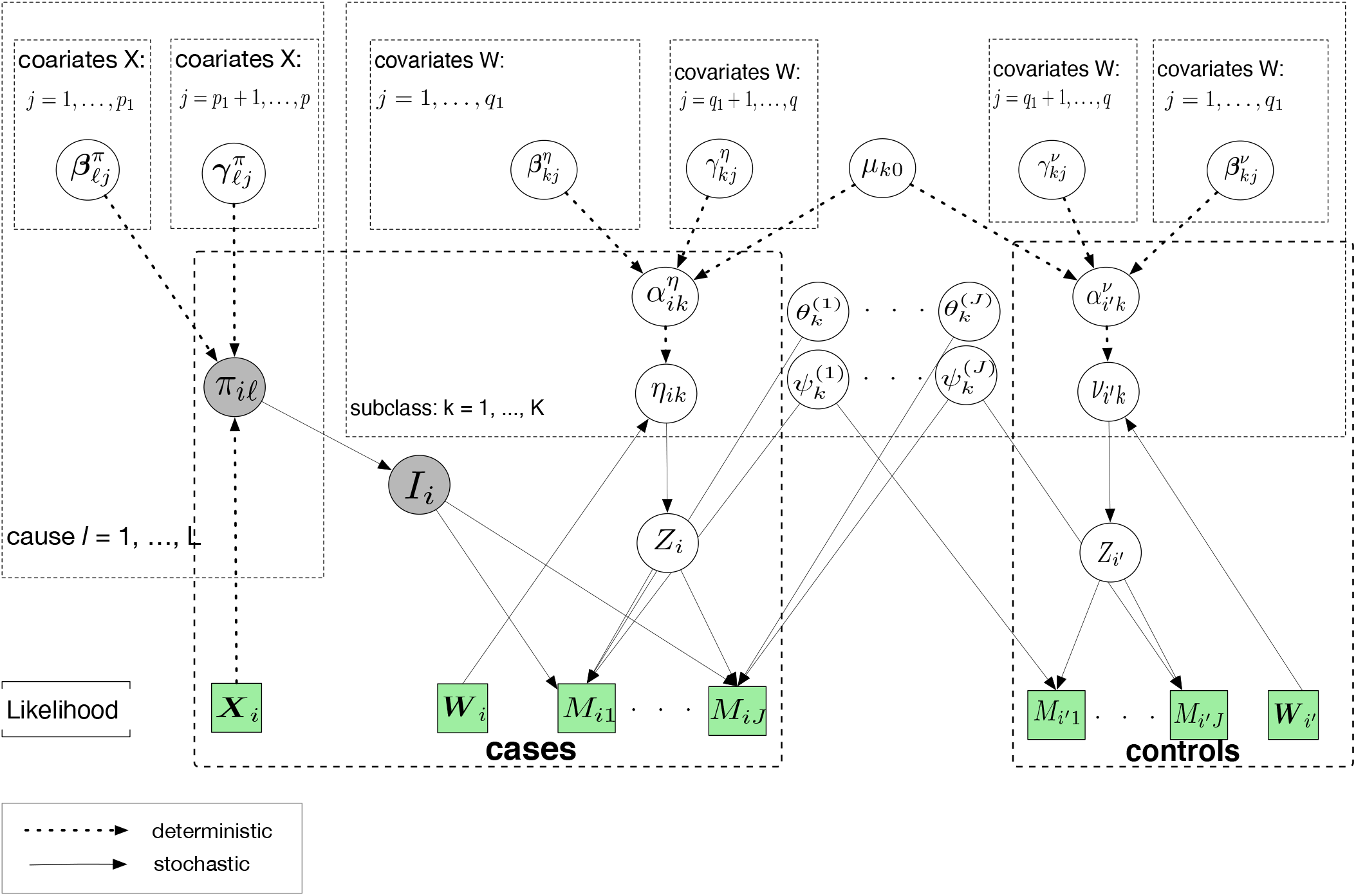
The directed acyclic graph (DAG) representing the structure of the model likelihood. The quantities in squares are either data or hyperparameters; the unknown quantities are shown in the circles. The arrows connecting variables indicate that the parent parameterizes the distribution of the child node (solid lines) or completely determines the value of the child node (dotted arrows). The rectangular “plates” where the variables are enclosed indicate that a similar graphical structure is repeated over the index; The index in a plate indicate subjects, causes, covariates or subclasses. Figure S6 in the Supplementary Materials presents the complete DAG with prior specification.

**FIGURE 2.**
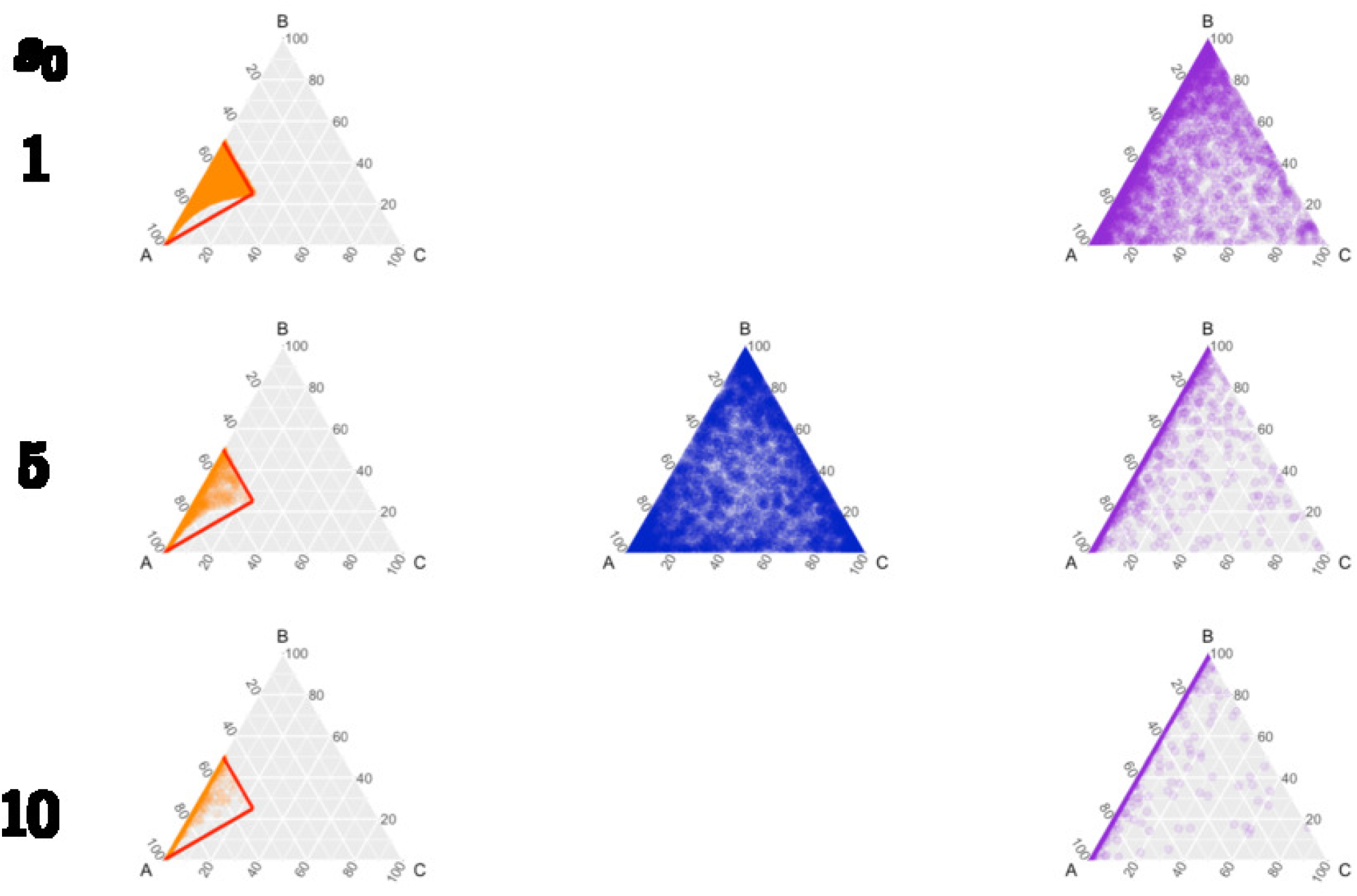
Random draws from the prior for three subclass weights (*ν_A_, ν_B_, ν_c_*)^⊤^ shown in ternary diagrams. The scale parameter *s*_0_ increases from 1 to 10 from the top to the bottom while fixing *ν*_0_ = 1. The random samples from the prior are increasingly concentrated towards the first subclass *A*. In each row, the three columns correspond to random samples when 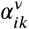 equals *μ*_*k*0_ in the left, 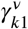 in the middle, and 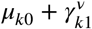 in the right. Here we show a single ternary diagram in the middle because the Gaussian prior of 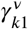 does not depend on *s*_0_: prior means are −1.07 and 0 for 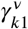, *k = A, B*, respectively, and the prior precision is *κ_γ_* = 1/4. These values were chosen to produce approximately evenly-distributed draws in a ternary diagram (middle).

#### 5.4.2 | Priors: Encourage Constant Additive Regression Functions: 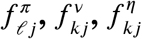

We use B-splines to approximate the additive functions of a standardized continuous variable^17^: 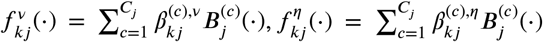, where 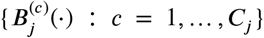 are *C_j_* cubic B-spline bases that are shared between the cases and the controls for the *j*-th covariate ***W**_j_* in subclass *k*. We assume distinct coefficients: 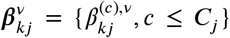 and 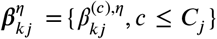. In addition, we assume 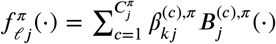 where 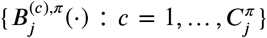 are also cubic B-spline bases and 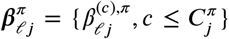. With *M* interior equally-spaced knots ***κ*** = (*κ*_0_,…, *κ*_*M*+1_)^⊤^. For example, for covariate *W_ij_* min_*i*_(*W_ij_*) = *κ*_0_ < *κ*_1_ < ⋯ < *κ_M_* < *κ*_*M*+1_ = max_*i*_(*W_ij_*), there are *M* + 4 basis functions. It readily extends to different numbers of basis functions. We further restrict 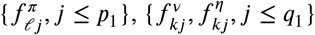 to have zero means for statistical identifiablity.

The choice of the number of bases (or, degrees of freedom) is crucial for the estimation of the CSCF functions and subclass weight regression functions. For example, larger values of *C_j_* and 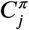 define a richer class of functions that can accommodate abrupt seasonal effect. In the context of PERCH study, more parsimonious models are preferred. Further improvements in knot selection is possible using knots on a nonequidistant grid so that more knots are placed where cases and controls are dense. Our practical suggestion is to place 5 to 10 equally spaced knots. A more computationally intensive approach is to add or delete knots using split-merge type of algorithms.^18^

Since the prior specifications below apply to 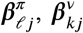 and 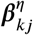, for notational simplicity, we omit the superscripts *π, ν, η* and use “•” as a placeholder. We specify Gaussian random walk priors on the basis coefficients via Bayesian P-splines.^17^ Let 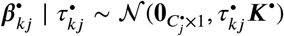, where the symmetric penalty matrix 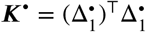 is constructed from the first-order difference matrix 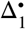 of dimension 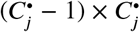. It maps adjacent B-spline coefficients to 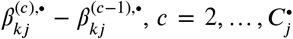. The precision matrix 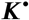 is not full rank and hence leaves the prior of 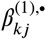 unspecified. We assume independent priors 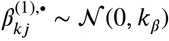.

Importantly, the 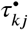 is the smoothing parameters. Large values of 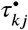 lead to smoother fit of 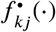 (constant when 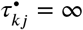).

We specify a mixture prior for smoothing parameter 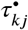:

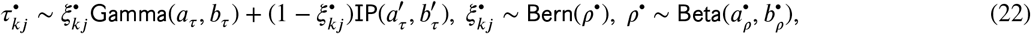

where the Gamma-distributed component concentrates near smaller values and the inverse-Pareto component 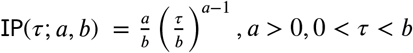, prefers larger values, and 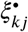. is a binary indicator for the Gamma component. This bimodal mixture distribution creates a sharp separation between flexible and constant fits.^19,20^

#### 5.4.3 | Prior Distributions for Other Parameters

We assume independent Gaussian priors 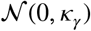 for each element of 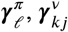 and 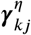. See Appendix A2 in the Supplementary Materials for the choice of hyperparameters *ν*_0_ and *s*_0_ in (21), 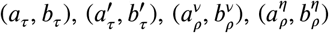 and 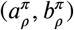 in (22), *κ_β_*, and *κ_γ_*. The npLCM regression model is partially-identified.^21^ We assume independent 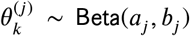, *j ≤ J*. In real data applications, hyperparameters (*a_j_,b_j_*) are chosen so that the 2.5% and 97.5% quantiles match an elicited prior range.^22^ For FPRs, we assume a flat prior 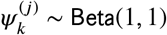 because they are empirically estimable from the control data.

### 5.5 | Posterior Inference and Software

Figure 1 summarizes the generative process above and the priors in a directed acyclic graph (DAG). Appendix A3 in the Supplementary Materials uses the DAG to derive the Markov chain Monte Carlo (MCMC) algorithm that draws posterior samples of the unknowns to approximate their joint posterior distribution.^23^ Flexible posterior inferences about any functions of the model parameters and individual latent variables are available by plugging in the posterior samples of the unknowns. All the models in this paper are fitted using a free and publicly available `R package baker` (https://github.com/zhenkewu/baker).

## 6 | SIMULATIONS

We simulate case-control non-gold-standard diagnostic test data along with observed continuous and/or discrete covariates under multiple combinations of true parameter values and sample sizes that mimic the motivating PERCH study. In Simulation I, we illustrate flexible statistical inferences about the CSCF functions. In Simulation II, we focus on the overall CSCFs ***π***\* in (14) with *G*(·) being the empirical distribution of {***X**_i_*}. We compare the frequentist properties of the posterior mean ***π***\* obtained from analyses with or without covariates upon repeated use across independent replications.^24^ We compare the proposed model with npLCMs without covariates, because the latter is the only available method for estimating CSCFs using case-control data. Regression analyses reduce estimation bias, retain efficiency and provide more valid frequentist coverage of the 95% credible intervals (CrIs). The relative advantage varies by the true data generating mechanism and sample sizes.

In all analyses below, to mimic weakly informative prior for TPRs, we use independent Beta(7.13,1.32) TPR prior distributions that match a wide interval 0.55 to 0.99 with the lower and upper 2.5% quantiles, respectively. We tested other priors that matched slightly wider or narrower ranges and observed similar advantages of the proposed model. The priors for other parameters are specified in Section 5.4.

### Simulation I

We demonstrate posterior inference of true CSCF functions 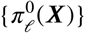. We let *π_ℓ_*(·), *ν_k_*(·) and *η_k_*(·) depend on the two covariates ***X = W*** = (*S, T*), *S* and enrollment date (*T*), so that regression adjustments are necessary (see Remark 2). We simulate *N_d_* = 500 cases and *N_u_* = 500 controls for each of two levels of *S* (a discrete covariate) and uniformly sample the subjects’ enrollment dates over a period of 300 days. Appendix A4 in the Supplementary Materials specifies the true data generating mechanism. Based on the simulated data, pathogen A has a bimodal case positive rate curve mimicking the trends observed of RSV in one PERCH site. Other pathogens have overall increasing case positive rate curves over enrollment dates. We set the simulation parameters in a way that the *marginal* control rate may be higher than cases for earlier enrollment dates. Row 2 of Figure 3 shows that for the 9 causes in the columns, the posterior means and 95% CrIs for the CSCF functions *π_ℓ_*(·) well recover the simulation truths 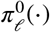 in the Stratum *S* = 1; similarly good recovery is observed for stratum *S* = 2. Figure S1 inthe Supplementary Materials further demonstrates well-recovered subclass weight curves. Appendix A5 in the Supplementary Materials provides additional simulation results that shows the true 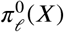 is well-recovered for a discrete covariate *X*.

**FIGURE 3.**
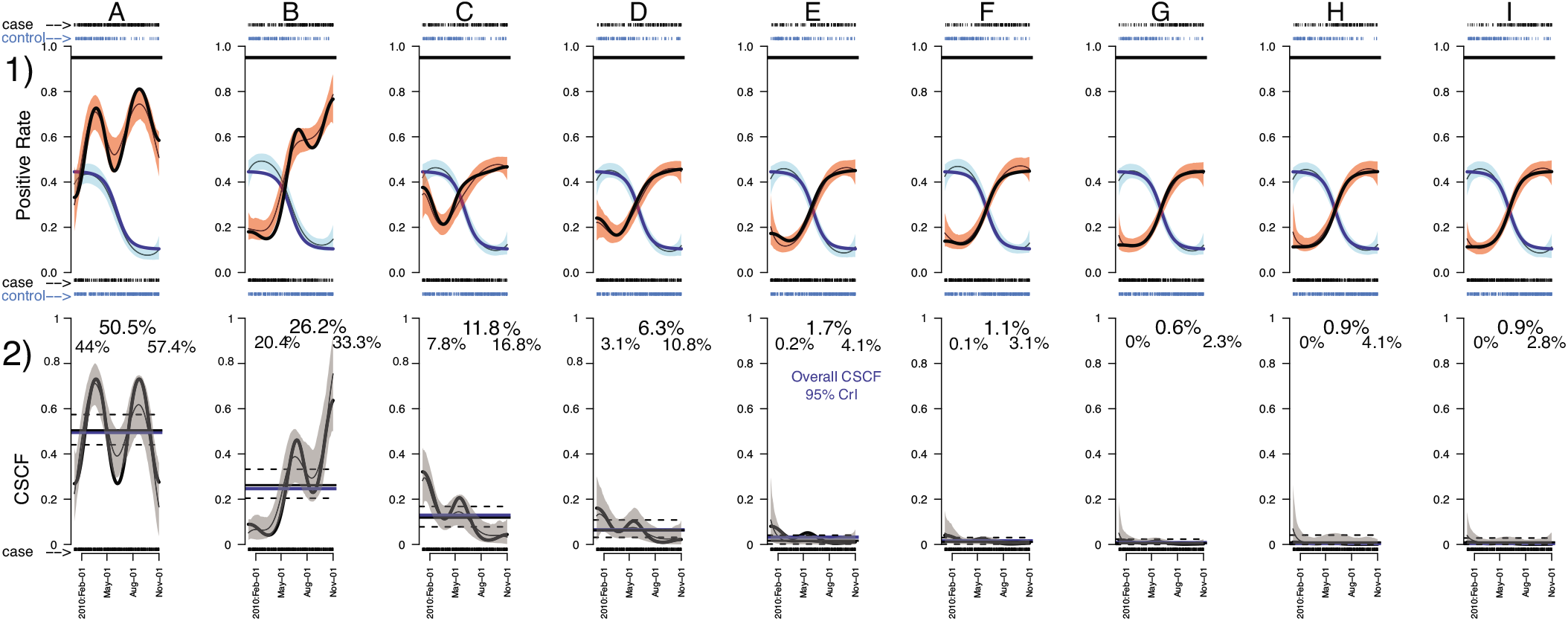
Row 2) For each of the 9 single-agent causes (by column) in Simulation I, the posterior mean (thin black curves) and pointwise 95% CrIs (gray) for the CSCF curves *π_ℓ_*(*x*) are close to the simulation truths 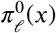 in the stratum of *S* = 1 (thick black curves). The overall CSCF in the particular stratum is shown along with the 95% CrI on both sides.^25^ In row 1), the fitted case (red, 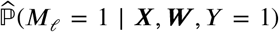) and control (blue, 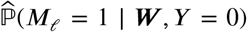) positive rate curves are shown with the posterior mean curves (thin black curves) along with pointwise 95% credible bands; The rug plots show the positive (top) and negative (bottom) measurements made on cases and controls on the enrollment dates. The solid horizontal lines in row 1 indicate the true TPRs.

### Simulation II

We show the regression model accounts for population stratification by covariates hence reduces the bias of the posterior mean 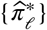 in estimating the overall CSCFs (***π***\*) and produces more valid 95% CrIs. We illustrate the advantage of the regression approach under simple scenarios with a single two-level covariate *X* ∈ {1, 2}. We set *W = X*. We perform npLCM regression analysis with *K*= 3 for each of *R* = 200 replication data sets simulated under each scenario detailed in Appendix A4 in the Supplementary Materials corresponding to distinct numbers of causes, sample sizes, relative sizes of CSCF functions (rare versus popular causes), signal strengths (more discrepant TPRs and FPRs indicate stronger signals), and effects of *W* on {*ν_k_*(*W*)} and {*η_k_*(*W*)}.

In estimating 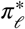, we focus on evaluating the marginal bias 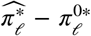, where 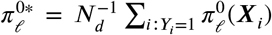 is the true overall CSCF, and 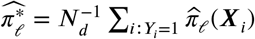 is an empirical average of the posterior mean CSCFs at ***X**_i_*. We also evaluate the empirical coverage rates of the 95% CrIs.

The proposed model incorporates covariates and performs better in estimating ***π***\* than a model that omits covariates. For example, Figure 4(a) shows for *J* = 6 that, relative to no-covariate npLCM analyses, regression analyses produce posterior means that on average have negligible relative biases (percent difference between the posterior mean and the truth relative to the truth) for each pathogen across simulation scenarios. As expected, we observe slight relative biases from the regression model in the bottom two rows of Figure 4(a), because the informative TPR prior Beta(7.13,1.32) has a mean value lower than the true TPR 0.95. A more informative prior centered closer to the true TPR would further reduce the relative bias. See additional simulations in Appendix A5 in the Supplementary Materials on the role of informative TPR priors. Regression analyses also produce 95% CrIs for 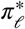 that have more valid empirical coverage rates in all the scenarios (Figure 4(b)). Misspecified models without covariates concentrate the posterior distribution away from the true overall CSCFs, resulting in severe under-coverage.

**FIGURE 4.**
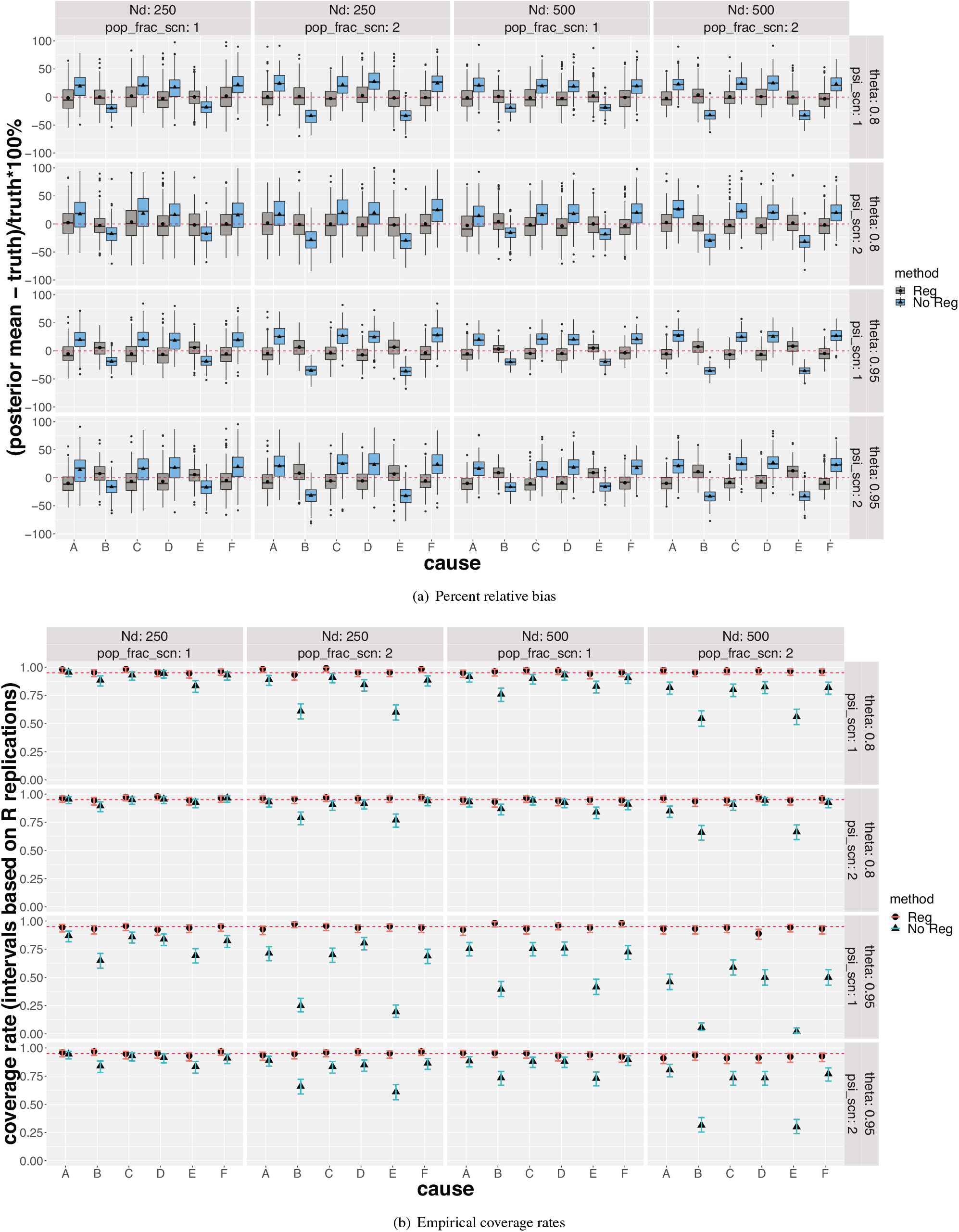
The regression analyses produce less biased posterior mean estimates and more valid empirical coverage rates for 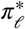 in Simulation II. Each panel corresponds to one of 16 combinations of true parameter values and sample sizes. *Top*) Each boxplot (left: regression; right: no regression) shows the distribution of the percent relative bias of the posterior mean in estimating the overall CSCF 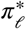 for six causes (A - F); 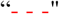 indicates zero bias. *Bottom*) The empirical coverage rates of the 95% CrIs with regression (•) or without regression (▲); 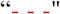 indicates the nominal 95% level. Since each coverage rate for 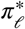 is computed from *R* = 200 binary observations of the true 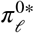 being covered or not, a 95% CI is also shown.

The proposed model is flexible in incorporating covariates which enables the borrowing of information from subjects with similar covariates. As expected, we observed improved prediction performance in simulation settings. As a more realistic test of the proposed method, we will discuss in more detail the comparison of the relative predictive performances of the proposed method against a closely related stratified analytic approach on the real data analysis in Section 7.

The hyperparameter choice is critical for adequate model performance, e.g., in terms of producing small posterior mean squared error (PMSE). We conducted sensitivity analysis for the hyperparameter that controls the sparseness of subclass weights at values *ν*_0_ = 0.01, 0.1, 1, 10, 100. We observed that the influence of hyperparameter upon the PMSE for CSCFs is stronger when the parameters settings correspond to weaker signals, e.g., when 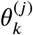 is closer to 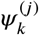; the results are stable for different hyperparameter choices under simulation scenarios corresponding to stronger signals.

## 7 | RESULTS FROM PERCH STUDY

We restrict attention in this regression analysis to 518 cases and 964 controls from one of the PERCH study sites in the Southern Hemisphere that collected more complete information on enrollment date (*t*, August 2011 to September 2013; standardized), age (dichotomized to younger or older than one year), HIV status (positive or negative), disease severity for cases (severe or very severe), and presence or absence of seven species of pathogens (five viruses and two bacteria, representing a subset of pathogens evaluated) in NPPCR.

The names of the pathogens and the abbreviations are (i) bacteria: *Haemophilus influenzae* (`HINF`) and *Streptococcus pneumoniae* (`PNEU`), (ii) viruses: adenovirus (`ADENO`), human metapneumovirus type A or B (`HMPV_A_B`), parainfluenza type 1 virus (`PARA_1`), rhinovirus (`RHINO`), and respiratory syncytial virus (`RSV`).

We also include in the analysis the case-only, perfectly specific but imperfectly sensitive blood culture (BCX) diagnostic test results for two bacteria from cases only. For BCX data, we assume perfect specificity which is guided by the fact that if a pathogen did not infect the lung, it cannot be cultured from the blood (so we do not need control data to estimate the specificities). Detailed analyses of the entire data are reported elsewhere.^1^ Table 1 shows the observed frequencies in the ***W*** = (`age, HIV status`) strata for controls and ***X*** = (`age, HIV status, disease severity`) strata for cases. The two case strata with the most subjects are severe pneumonia children who were HIV negative, under or above one year of age. Table 1 has small cell counts.

**TABLE 1.** The observed counts (frequencies) of controls by age and HIV status; Case counts are further stratified by disease severity (1: yes; 0: no). The observed marginal rates are shown at the bottom. Enrollment date (*t*) is not stratified upon here.

| age $\geq 1$ | HIV positive | # controls (%)<br>total: 964 (100) | very severe (VS)<br>(case-only) | # cases (%)<br>total: 518 (100) |
| --- | --- | --- | --- | --- |
| 0 | 0 | 548 (56.8) | 0<br>1 | 208 (40.2)<br>120 (23.2) |
| 1 | 0 | 280 (29.0) | 0<br>1 | 69 (13.3)<br>32 (6.2) |
| 0 | 1 | 85 (8.8) | 0<br>1 | 37 (7.1)<br>25 (4.8) |
| 1 | 1 | 51 (5.3) | 0<br>1 | 24 (4.6)<br>3 (0.6) |
| case: 24.7% | 17.2% |  | 34.7% |  |
| control: 34.3% | 14.1% |  | - |  |

Regression modeling techniques must be used to deal with small strata of cases and controls and obtain uncertainty quantification. Consider the discrete covariates only: a naive *fully-stratified* analysis that fits an npLCM to the case-control data in each covariate stratum is problematic. First, sparsely-populated strata defined by many discrete covariates may lead to unstable CSCF estimates. Second, it is often of policy interest to estimate the overall CSCFs ***π***\*. Since the informative TPR priors are often elicited for a case population and rarely for each stratum, reusing independent prior distributions of the TPRs across all the strata during multiple npLCM fits will lead to overly-optimistic posterior uncertainty in ***π***\*, hampering policy decisions. Third, relative to controls, cases may be further stratified by additional covariates (e.g., disease severity), resulting in finer case strata nested in each control stratum (e.g., see Table 1). Because every npLCM fit requires a case and control sample, a control stratum would have to be reused for every finer case strata. We use the proposed regression model to address these issues.

Figure S4 in the Supplementary Materials shows summary statistics for the NPPCR and BCX data including the positive rates in the cases and the controls and the conditional odds ratio (COR) contrasting the case and control rates adjusting for the presence or absence of other pathogens (NPPCR data only).

To fit the model, we include in the regression analysis seven single-pathogen causes 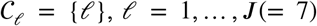 and a “`Not Specified (NoS)`” cause denoted by 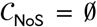 to account for other non-targeted causative agents. We incorporate the prior knowledge about the TPRs of the NPPCR measures from laboratory experts. We set the Beta priors for sensitivities by *a_θ_* = 12.68 and *b_θ_* = 4.83, so that the 2.5% and 97.5% quantiles match the lower and upper ranges of plausible sensitivity values of 0.5 and 0.9, respectively. We use a working number of subclasses *K* = 5. Results under larger *K*s remain nearly the same. In the presence of BCX data for a subset of two bacteria (only bacteria can be cultured from blood), because BCX data have very low prevalence, we multiple *L*^reg^ by BCX data likelihood in the simpler pLCM (2)-(4) and specify the `Beta`(7.59,58.97) prior for the two TPRs of BCX measurements matching the range of 5 - 20% based on existing vaccine probe trials.^26^ In the CSCF regression model 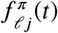, we use 7 d.f. for B-spline expansion of the additive function for the standardized enrollment date *t* at uniform knots along with three binary indicators for age older than one, HIV positive, very severe pneumonia. In the subclass weight regression model, we use 5 d.f. for the standardized enrollment date *t* with uniform knots and two indicators: **1**{`age`_*i*_ ≥ 1 `year`} and **1**{`HIV`_*i*_ = 1}. The prior distributions for other parameters follow the specification in Section 5.4.

The regression analysis produces seasonal estimates of the CSCF function for each cause that varies in trend and magnitude among the eight case strata defined by age, HIV status and disease severity. Figure 5 shows for two age-HIV-severity strata the posterior mean CSCF curves and 95% pointwise credible bands of the *π_ℓ_*(*t*, `age, HIV, severity`) as a function of *t*. For example, among the younger, HIV negative and severe pneumonia children (Figure 5(a)), the CSCF curve of `RSV` is estimated to have a prominent bimodal temporal pattern that peaked at two consecutive winters in the Southern Hemisphere (June 2012 and 2013), suggesting prioritization of preventative measures and treatment algorithms for RSV. Other single-pathogen causes `HINF, PNEU, ADENO, HMPV_A_B` and `PARA_1` have overall low and stable CSCF curves across seasons. As a result, the estimated CSCF curve of `NoS` shows a trend with a higher level of uncertainty that is complementary to `RSV`. In contrast, Figure 5(b) shows a lower degree of seasonal variation of the `RSV` CSCF curve among the older, HIV negative and severe pneumonia children.

**FIGURE 5.**
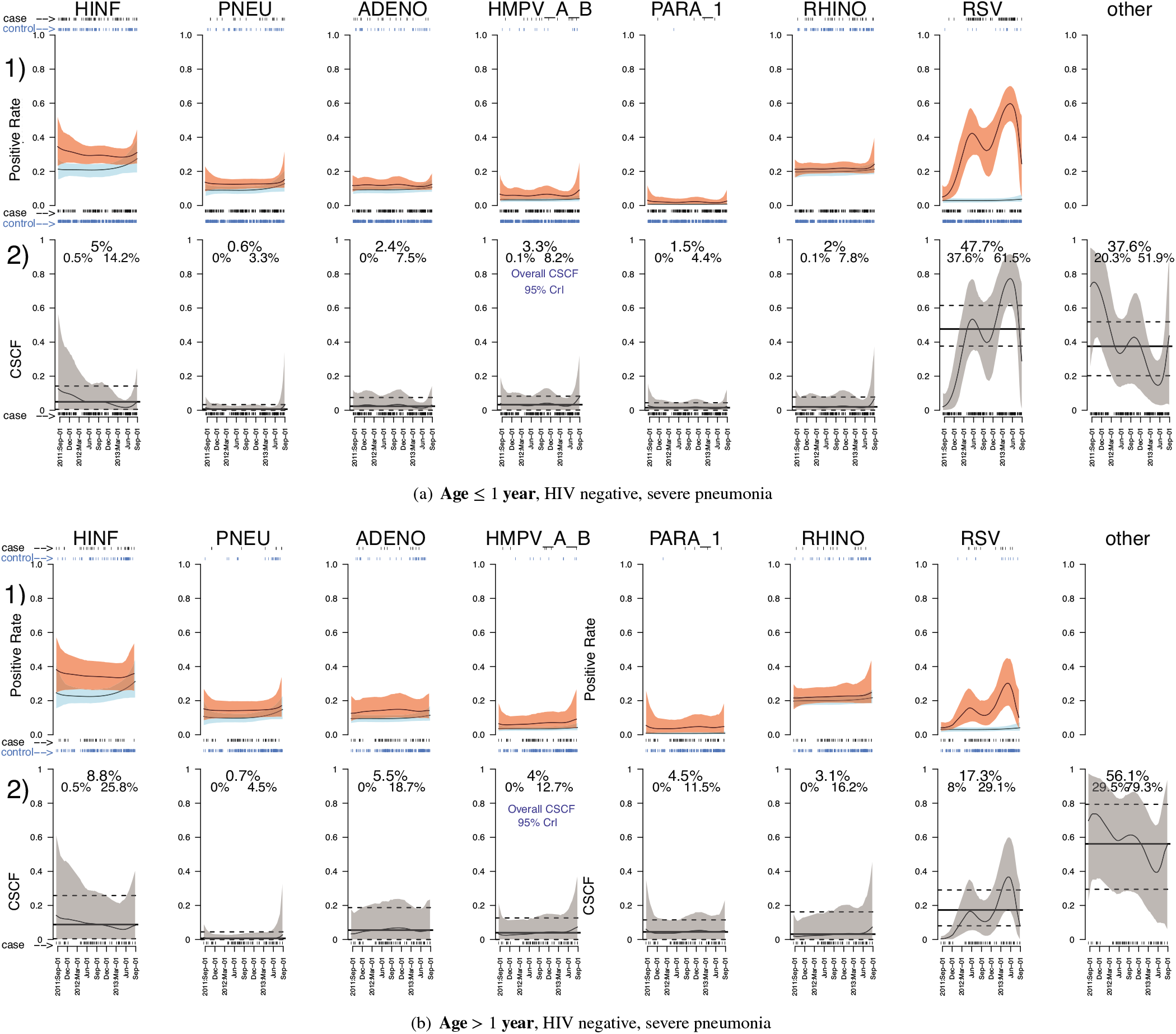
Estimated seasonal CSCF for two most prevalent age-HIV-severity case strata under single-pathogen causes (`HINF, PNEU, ADENO, HMPVAB, PARA1, RHINO, RSV`) and a “Not Specified” cause. In each age-HIV-severity case stratum and for each cause *ℓ*: Row 2): Temporal trend 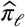(`t; age, HIV, severity`) enveloped by pointwise 95% CrIs (gray). The horizontal solid line shows the estimated overall CSCF 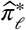(`age, HIV, severity`) averaged over cases in the present stratum (dashed black lines: 95% CrI). The rug plot on the x-axis indicates cases’ enrollment dates. Row 1) shows the fitted temporal case (red) and control (blue) positive rate curves enclosed by the pointwise 95% CrIs; The two rug plots at the top (bottom) indicate the dates of the cases and controls being enrolled and tested positive (negative) for the pathogen.

The inferential algorithm based on the regression model can also perform individual-level cause-specific probability assignment given a case’s measurements and automatically use covariate values during assignment. Figure S5 in the Supplementary Materials show distinct cause-specific probabilities for two cases (one older than one and the other younger than one) with all-negative NPPCR results.

We estimate the overall CSCFs ***π***\*(`age, HIV, severity`) in every age-HIV-severity stratum by averaging the CSCF function estimates over the empirical distribution of enrollment date in each stratum. For example, contrasting the two age-HIV-severity strata in Figure 5(a) and 5(b), the case positive rate of `RSV` among the older children drops from 39.3% to 17.9% but the control positive rates remain similar (from 3.0% to 4.1%). The overall CSCF estimate for `RSV` 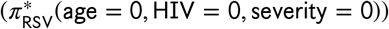 drops from 47.7 (95% CrI: 37.6,61.5)% to 17.3 (95% CrI: 8.0,29.1)%. The CSCF estimate for `NoS` 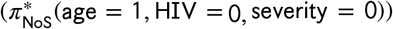 increases from 37.6 (95% CrI: 20.3,51.9)% to 56.1 (95% CrI: 29.5,79.3)%. The overall CSCFs for other causes remain similar between the younger and older, HIV negative severe pneumonia children.

### Comparison with Other Methods

The most relevant comparison method is based on stratum-specific npLCM analysis. Because in the data analysis, the enrollment date is continuous, stratification requires categorization of the enrollment date into “time periods”, which after discussion with scientists in PERCH study is not straightforward. We decided to only stratify on the three binary covariates used in the main analysis, `I`(`age > 1`), `I`(`HIV positive`), `I`(`very severe pneumonia`). Because in the PERCH study, case and control enrollment are frequency matched, leaving out enrollment date still provides a reasonable comparison against the main method. We re-emphasize that, however, the stratified analysis cannot output a CSCF as a function of continuous covariates such as enrollment date.

We compare the proposed and the stratum-specific analysis via prediction of the multivariate binary responses {***M**_i_*} from the diagnostic tests. We define the prediction based on the posterior mean of the positive response probabilities given all the data 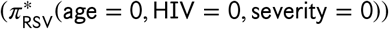. We predict a response *M_ij_* to be 1 if the posterior mean of the positive response probability given the individual’s covariate values exceeds 0.5, i.e. 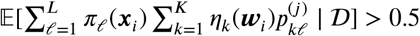; otherwise, we let the prediction be 0. The prediction error is defined as the fraction of incorrectly predicted responses. We found that the proposed regression method has gained additional ability to predict the observed responses compared to the stratum specific method. This is partly due to the ability of the regression model to borrow information across subjects with different covariate values, e.g., through additive assumptions and smoothing over the continuous enrollment date. We also conducted a no-covariate analysis and compared its prediction performance; the complete no-covariate analysis results are shown in Figure S4 in the Supplementary Materials. Among the three methods, proposed regression model, stratified analysis and no-covariate analysis, the prediction errors are 10.3%, 17.8%, 24.7%, respectively, with the smallest prediction error based on the proposed regression model.

## 8 | DISCUSSION

In disease etiology studies where gold-standard data are infeasible to obtain, epidemiologists need to integrate case and control data to draw inference about the population CSCFs and individual cause-specific probabilities that may depend on covariates. The only existing methods for case-control data based on npLCM do not describe the relationship between covariates and the CSCFs. This paper fills this analytic gap by extending npLCM to a unified hierarchical Bayesian regression modeling framework. The model-based inferential algorithm estimates CSCF functions and is amenable to efficient posterior computation.

We have shown via simulations that the regression approach accounts for population stratification by important covariates and, as expected, reduces estimation biases and produces 95% credible intervals that have more valid empirical coverage rates than an npLCM analysis that omits covariates. In addition, the proposed regression analysis can readily integrate multiple sources of diagnostic measurements with distinct levels of diagnostic sensitivities and specificities, a subset of which are only available from cases. Our regression analysis uses data from one PERCH site with more complete covariate information and reveals prominent dependence of the CSCFs of the virus RSV upon seasonality and a pneumonia child’s age, HIV status and disease severity.

The proposed approach has three distinguishing features: 1) It allows an analyst to specify a model for the functional dependence of the CSCFs upon important covariates. Assumptions such as additivity further improves estimation stability for sparsely-populated strata defined by many discrete covariates. 2) The model incorporates control data to infer the CSCF functions. The posterior algorithm estimates a parsimonious covariate-dependent reference distribution of the diagnostic measurements from controls, which is critical for correctly assigning cause-specific probabilities for individual cases. Finally, 3) the model uses informative priors of the sensitivities (TPRs) only once in the entire target population for which these priors were intended. Relative to a fully-stratified npLCM analysis that reuses these TPR priors, the proposed regression analysis avoids overly-optimistic uncertainty estimates for the overall CSCFs.

The covariate-stratified approach have the advantage of being simple-to-understand and easy to implement given the previous generation of the nested partially-latent class model. However, the pain points are 1) its requirement on the sample size for each covariate stratum, and 2) the controls may be reused as statistical comparison group for multiple case stratum. As shown in Table 1, both issues appeared. First, small sample sizes in many strata (e.g., the final row has only three cases: age older than 1, HIV positive, very severe) result in a stratified analysis that is not statistically stable. Turning to the second issue of selecting the control group, a naive stratified analysis would need to use the 51 controls twice, one for the not very severe case group (24 cases) and the other for very severe case group (3 cases). Reusing control data twice may lead to incorrect variance estimate of the CSCFs. The above two issues prevented producing reliable results based on the stratified analysis. However, we provided comparison results with only discrete covariates in the regression framework and showed advantage of the proposed regression approach that can handle both continuous and discrete covariates.

Future work may further expand the utility of the proposed methods. First, flexible and parsimonious alternatives to the additive models may capture important interaction effects.^27^ Second, in the presence of many covariates, class-specific predictor selection methods for *π_ℓ_*(***X**_i_*) may provide further regularization and improve interpretability^28^. Third, when the subsets of pathogens 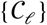 that have caused the diseases in the population are unknown, the proposed method can be combined with subset selection procedures^29^. Finally, wide applicability of posterior sampling algorithms have increased the use of complex and multilevel hierarchical models like the proposed model in this paper. Other methods for hyperparameter choice may be marginal likelihood maximization, specifying another layer of hyperpriors, or empirical Bayes approach. A more thorough comparative study for different methods of hyperparameter choice is warranted. We leave this for future work.

## Supporting information

The Supplementary Materials contain the technical details, extra simulation results and figures referenced in Main Paper.

## ACKNOWLEDGMENTS

We thank the PERCH study team led by Katherine O’Brien for providing the data and scientific advice, Scott Zeger, Maria Deloria-Knoll, Christine Prosperi and Qiyuan Shi for valuable feedback about `baker` and Jing Chu for preliminary simulations. We are grateful for comments from the Editor, Associate Editor and two reviewers that greatly improved the presentation.

## SUPPORTING INFORMATION

Additional supporting information may be found online in the Supporting Information section at the end of this article.

### How to cite this article

Z. Wu and I. Chen (2020+), Probabilistic Cause-of-disease Assignment using Case-control Diagnostic Tests: A Hierarchical Bayesian Latent Variable Regression Approach, *Statistics in Medicine, 20XX;XX:X–X*.

