## Supplementary material for "Probabilistic Cause-of-disease Assignment using Case-control Diagnostic Tests: A Latent Variable Regression Approach": The Supplementary Materials contain the technical details, extra simulation results and figures referenced in Main Paper.

### Supplementary Materials for "Probabilistic Cause-of-disease Assignment using Case-control Diagnostic Tests: A Hierarchical Bayesian Latent Variable Regression Approach"

ZHENKE WU[*,1,2] and IRENA CHEN[1]

[1] *Department of Biostatistics, University of Michigan, Ann Arbor, MI 48109, USA*

[2] *Michigan Institute for Data Science, University of Michigan, Ann Arbor, MI 48109, USA*

[*] **

SUMMARY

The Supplementary Materials present the technical details, additional extensive simulation results and supplemental figures referenced in the Main Paper. Section A1 remarks on the assumptions for the control model. Section A2 discusses the choice of hyperparameters. Section A3 provides the posterior algorithm and convergence checks for valid posterior inference. Section A4 describes the simulation settings in the Main Paper and Section A5 shows additional simulation results. Section A6 presents additional results from PERCH study. Section A7 contains Supplemental Figures.

Motivated by the data application in PERCH study, we will use some existing terminologies (Wu *and others*, 2016) for two sources of imperfect diagnostic tests of the pathogen causes (those infecting the lung): (a) case-control tests: NP PCR results that are not perfectly sensitive or specific, referred to as "bronze-standard" (BrS) data; and (b) case-only tests: blood culture (BCX) results for a subset of pathogens that are perfectly specific but lack sensitivity, referred to as "silver-standard" (SS) data.

[*]To whom correspondence should be addressed.

#### A1. Remark on the Assumption for Introducing Covariates into the Control Model

By assuming mutually independent measurements $M_1, \ldots, M_J$ given subclass $Z$ and $Y = 0$, we let the covariates influence the dependence structure of the measurements only through the unobserved $Z$. As a result, upon integrating over $Z$, the proposed model does *not* assume marginal independence $\mathbb{P}(\boldsymbol{M} \mid \boldsymbol{W}, Y = 0) = \prod_{j=1}^{J} \mathbb{P}(M_j \mid \boldsymbol{W}, Y = 0)$ in contrast to a kernel-based method that makes this assumption (Saha *and others*, 2018, Supplementary appendix). Our approach to incorporating covariates to model control data follows the latent class regression formulation in Bandeen-Roche *and others* (1997). This formulation is classical and has been used in applications including HIV population size estimation (Bartolucci and Forcina, 2006), and alcoholic and drug addiction (Chung, Flaherty and Schafer, 2006).

#### A2. Considerations for the Choice of Hyperparameters

In this paper, we use $(a_\tau, b_\tau) = (3, 2)$ and $(a'_\tau, b'_\tau) = (1.5, 400)$ in the Gamma-Inverse-Pareto mixture prior for the precision parameter in the Gaussian random walk prior. We use the same Beta hyperparameters for $\rho^\nu$ and $\rho^\eta$ ($a_\rho = 0.5, b_\rho = 1$) in the control and case subclass regressions to *a priori* give slight preference for constant curves; We use $a^\pi_\rho = 1$, $b^\pi_\rho = 0.5$ to *a priori* give slight preference for flexible CSCF functions.

In our simulations and applications, we choose hyperparameters $\nu_0 = 1$ and $s_0 = 10$ for the intercept, and $k_\beta = 1/4$ for the first B-spline coefficients $\beta^{(1),\nu}_{kj}$, $\beta^{(1),\eta}_{kj}$ in the random walk prior in P-spline. We have chosen our hyperparameters based on the interpretations on the probability (inverse-link) scale; see similar prior elicitations for regression coefficients in other applications (e.g., Bedrick, Christensen and Johnson, 1996; Witte, Greenland and Kim, 1998) and for automatic, stabilized and weakly-informative fitting of generalized linear models (Gelman *and others*, 2008). We choose the hyperparameters for the intercepts that put most prior mass of $g(\mu_{10})$ within $(0.5, 1-10^{-9})$, because $1-10^{-9}$ is sufficiently close to 1 which means the stick-breaking is stopped at Step $k=1$. In contrast, we choose the first B-spline coefficient's hyperparameter $k_\beta = 1/4$ that puts most prior mass of $g(\beta_{kj}^{(1),\nu})$ within $(0.02, 0.98)$, a range for the weight of a non-trivial subclass to break from the rest of the stick at Step $k$. Note the first B-spline coefficient $\beta_{\ell j}^{(1),\pi}$ also has a $\mathcal{N}(0,\kappa_\beta)$ prior; Finally, we choose $\kappa_\gamma = \kappa_\beta = 1/4$.

#### A3. Posterior Sampling Algorithm

We update the unknowns by iterating through the following steps.

1. Update the class indicator $I_i \in \{1,\ldots,L\}$ for every case, from a categorical distribution with probability

$$\mathbb{P}(I_i = \ell \mid \mathit{others}) \propto [\boldsymbol{M}_i \mid I_i = \ell, Z_i, \boldsymbol{\Theta}, \boldsymbol{\Psi}][I_i = \ell \mid \pi_{i\ell}]$$
$$\propto \prod_{j \in \mathcal{C}_\ell} \left\{\theta_{Z_i}^{(j)}\right\}^{M_{ij}} \left\{1-\theta_{Z_i}^{(j)}\right\}^{1-M_{ij}} \cdot \prod_{j' \notin \mathcal{C}_\ell} \left\{\psi_{Z_i}^{(j')}\right\}^{M_{ij'}} \left\{1-\psi_{Z_i}^{(j')}\right\}^{1-M_{ij'}} \cdot \pi_{i\ell},$$

where $\mathcal{C} \subset \{1,\ldots,J\}$ represents the subset of causative agents for cause $\ell$.

2. Update the subclass indicator $Z_i \in \{1,\ldots,K\}$ for every case from a categorical distribution with probability

$$\mathbb{P}(Z_i = k \mid \mathit{others}) \propto [\boldsymbol{M}_i \mid Z_i = k, I_i, \boldsymbol{\Theta}, \boldsymbol{\Psi}][Z_i = k \mid \eta_{ik}, I_i]$$
$$\propto \prod_{j \in \mathcal{C}_{I_i}} \left\{\theta_{k}^{(j)}\right\}^{M_{ij}} \left\{1-\theta_{k}^{(j)}\right\}^{1-M_{ij}} \cdot \prod_{j' \notin \mathcal{C}_{I_i}} \left\{\psi_{k}^{(j')}\right\}^{M_{ij'}} \left\{1-\psi_{k}^{(j')}\right\}^{1-M_{ij'}} \cdot \eta_{ik}.$$

Update the subclass indicator $Z_{i'} \in \{1,\ldots,K\}$ for every control subjects from a categorical distribution with probability $\mathbb{P}(Z_{i'} = k \mid \mathit{others}) \propto [\boldsymbol{M}_{i'} \mid Z_{i'} = k, \boldsymbol{\Psi}][Z_{i'} = k \mid \nu_{i'k}] \propto \prod_{j=1}^J \left\{\psi_k^{(j)}\right\}^{M_{i'j}} \left\{1-\psi_k^{(j)}\right\}^{1-M_{i'j}} \cdot \nu_{i'k}$.

3. Update the control subclass regression coefficients $\mathbf{\Gamma}^\nu = [(\Gamma_1^\nu)^\top, \ldots, (\Gamma_{K-1}^\nu)^\top]^\top$ by

$$[\mathbf{\Gamma}^\nu \mid others] \propto \prod_{i':Y_{i'}=0} [Z_{i'} \mid \mathbf{\Gamma}^\nu, \{\mu_{k0}^*\}, \boldsymbol{W}_{i'}] \prod_{k=1}^{K-1} [\Gamma_k^\nu \mid \boldsymbol{b}^\nu, \mathbf{K}_k^\nu]$$
$$= \prod_{i':Y_{i'}=0} \prod_{k=1}^{K-1} \frac{\left\{\exp(\mu_{k0} + \mathcal{W}_{i'}^\top \Gamma_k^\nu)\right\}^{\mathbf{1}\{Z_{i'}=k\}}}{\left\{1 + \exp(\mu_{k0} + \mathcal{W}_{i'}^\top \Gamma_k^\nu)\right\}^{\mathbf{1}\{Z_{i'} \geqslant k\}}} \cdot \mathcal{N}(\Gamma_k^\nu; \boldsymbol{b}^\nu, \mathbf{K}_k^\nu),$$

where $\boldsymbol{b}^\nu = \mathbf{0}$ is the prior mean and

$$\mathbf{K}_k^\nu = \text{blkdiag}\left\{\left\{\tau_{kj}^\gamma \Delta_1^\top \Delta_1 + \kappa_\beta (1, \mathbf{0}_{(C_j-1)\times 1})^\top (1, \mathbf{0}_{(C_j-1)\times 1}), j = 1, \ldots, q_1\right\}, \underbrace{\kappa_\gamma, \ldots, \kappa_\gamma}_{q-q_1}\right\}$$

is the prior precision with $\Delta_1$ being the first-order difference matrix (Section 4.3.2 in the Main Paper) and $\mathbf{1}\{\mathsf{A}\} = 1$ if the statement $\mathsf{A}$ is true; 0 otherwise. The conditional density admits direct sampling via auxiliary Pólya-Gamma (PG) variables (Polson, Scott and Windle, 2013; Linderman, Johnson and Adams, 2015),

$$[\omega_{i'k}^\nu \mid others] \overset{d}{\sim} \mathsf{PG}(1, \alpha_{ik}^\nu = \mu_{k0} + \mathcal{W}_{i'}^\top \Gamma_k^\nu), \text{ for } k \leqslant Z_{i'},$$
$$[\Gamma_k^\nu \mid others] \propto \mathcal{N}(\mathbf{m}_k^\nu, (V_k^\nu)^{-1}),$$
$$V_k^\nu = \left\{\boldsymbol{\mathcal{W}}^\top \Omega_k^\nu \boldsymbol{\mathcal{W}} + \mathbf{K}_k^\nu\right\}^{-1},$$
$$\mathbf{m}_k^\nu = V_k^\nu (\boldsymbol{\mathcal{W}}^\top \Upsilon_k + \mathbf{K}_k^\nu \boldsymbol{b}^\nu),$$

for $k = 1, \ldots, \max_{i'} Z_{i'}$, where $\boldsymbol{\mathcal{W}}$ represents the design matrix for the control subclass weight regression where row $i'$ is $\mathcal{W}_{i'} = [\boldsymbol{B}_1^\top(W_{i'1}), \ldots, \boldsymbol{B}_{q_1}^\top(W_{i'q_1}), \widetilde{W}_{i',q_1+1}, \ldots, \widetilde{W}_{i'q}]^\top$, $\Omega_k^\nu = \text{diag}\{\omega_{i'k}^\nu \mathbf{1}\{Z_i \geqslant k\}\}$, $\Upsilon_k = \text{diag}\left\{(\mathbf{1}\{Z_{i'} = k\} - \frac{1}{2} - \mu_{k0})\mathbf{1}\{Z_{i'} \geqslant k\} : Y_{i'} = 0\right\}$; We update the PG auxiliary variables $\{\omega_{i'k}^\nu\}$ in parallel across controls and subclasses.

We similarly update the regression coefficients $\mathbf{\Gamma}^\eta$ in the case subclass regression by replacing the control design matrix $\boldsymbol{\mathcal{W}}$ with the counterpart among cases; Let $\mathcal{W}_i$ indicate the row in the case design matrix for case $i$.

4. Update $\mu_{k0}^*$ according to the full conditional distribution:

$$[\mu_{k0}^* \mid others] \propto \prod_{i:Y_i=1, Z_i \geqslant k} [Z_i \mid \{\mu_{h0}^*, \Gamma_h^\eta, h \leqslant k\}] \prod_{i':Y_{i'}=0, Z_{i'} \geqslant k} [Z_{i'} \mid \{\mu_{h0}^*, \Gamma_h^\nu, h \leqslant k\}] \cdot [\mu_{k0}^* \mid b^*, \tau_{k0}]$$

$$\propto \mathcal{N}_+(\mu_{k0}^*; b^*, \tau_{k0}) \cdot \prod_{k'=k}^{K-1} \left[ \prod_{i:Y_i=1} \frac{\{\exp(\alpha_{ik'}^\eta)\}^{\mathbf{1}\{Z_i=k'\}}}{\{1+\exp(\alpha_{ik'}^\eta)\}^{\mathbf{1}\{Z_i \geqslant k'\}}} \cdot \prod_{i':Y_{i'}=0} \frac{\{\exp(\alpha_{i'k'}^\nu)\}^{\mathbf{1}\{Z_{i'}=k'\}}}{\{1+\exp(\alpha_{i'k'}^\nu)\}^{\mathbf{1}\{Z_{i'} \geqslant k'\}}} \right],$$

where $b^* = 0$ is the prior mean, $\tau_{k0}$ is the prior precision, and $\alpha_{ik'}^\eta = \sum_{h=1}^{k'} u_{k'h}\mu_{h0}^* + \mathcal{W}_i^\top \{\Gamma_h^\eta Y_i + \Gamma_h^\nu(1-Y_i)\}$ and $\alpha_{i'k'}^\nu = \sum_{h=1}^{k'} u_{k'h}\mu_{h0}^* + \mathcal{W}_{i'}^\top \mathbf{\Gamma}_h^\nu$ are the linear predictors at the $k$-th stick-breaking events. We introduce auxiliary Pólya-Gamma variables for exact sampling:

$$[\omega_{ik}^* \mid others] \overset{d}{\sim} \mathsf{PG}\left(1, \alpha_{ik}^\eta Y_i + \alpha_{ik}^\nu (1-Y_i)\right), \text{ for a case or control } i, \text{ for } k \leqslant Z_i,$$

$$[\mu_{k0}^* \mid others] \overset{d}{\sim} \mathcal{N}_+(m_k^*, \tau_{k0}),$$

$$v_{k0} = \left\{ \sum_{i:\text{ all subjects}} \sum_{h=k}^{Z_i} \omega_{ih}^* + \tau_{0k} \right\}^{-1},$$

$$m_k^* = v_{k0} \left\{ \sum_{i:\text{ all subjects}} \sum_{h=k}^{Z_i} \left( s_{ih} - \omega_{ih}^* \left[ \underbrace{\sum_{j \leqslant h} u_{jh}\mu_{j0}^*}_{\mu_{h0}} + \mathcal{W}_i^\top \{\mathbf{\Gamma}_h^\eta Y_i + \mathbf{\Gamma}_h^\nu (1-Y_i)\} \right] \right) + \tau_{k0} b^* \right\},$$

for $k = 1, \ldots, \max_i Z_i$, where $s_{ih} = \mathbf{1}\{Z_i = h\} - \frac{1}{2}\mathbf{1}\{Z_i \geqslant h\}$.

5. Update the smoothing parameters $\tau_{kj}^\nu$ and smoothness selection indicator $\xi_{kj}^\nu$. First randomly switch $\xi_{kj}^\nu$ to $\xi_{kj}^{\nu*}$ either from 0 to 1, or 1 to 0 for $k = 1, \ldots, K-1$, $j = 1, \ldots, p_1$. Given the parameter $\tau_{kj}^\nu$, we propose its candidate $\tau_{kj}^{\nu*}$ from the log-normal distribution with log-mean parameter $\tau_{kj}^\nu$. We accept $(\tau_{kj}^{\nu*}, \xi_{kj}^{\nu*})$ with probability

$$\min\left\{1, \frac{p(\boldsymbol{\beta}_{kj}^\nu; \tau_{kj}^{\nu*})\pi(\tau_{kj}^{\nu*} \mid \xi_{kj}^{\nu*}) q(\tau_{kj}^\nu \mid \tau_{kj}^{\nu*}) q(\xi_{kj}^\nu \mid \xi_{kj}^{\nu*})}{p(\boldsymbol{\beta}_{kj}^\nu; \tau_{kj}^\nu)\pi(\tau_{kj}^\nu \mid \xi_{kj}^\nu) q(\tau_{kj}^{\nu*} \mid \tau_{kj}^\nu) q(\xi_{kj}^{\nu*} \mid \xi_{kj}^\nu)}\right\},$$

where $p(\boldsymbol{\beta}_{kj}^\nu; \tau_{kj}^{\nu*})$ is the density function $\mathcal{N}(0, \tau_{kj}^\nu \Delta_1^\top \Delta_1)$ and $\pi(\tau_{kj}^\nu \mid \xi_{kj}^\nu)$ is the prior distribution ((24) in the Main Paper).

We update $\tau_{kj}^{\nu}$ again because it is continuous and therefore has a much bigger parameter space than that of discrete parameter. Using random walk Metroplis-within-Gibbs algorithm, we propose $\tau_{kj}^{\nu *}$ from the log-normal distribution with log-mean parameter $\tau_{kj}^{\nu}$ and accept with probability

$$\min\left\{1, \frac{p(\boldsymbol{\beta}_{kj}^{\nu};\tau_{kj}^{\nu *})\pi(\tau_{kj}^{\nu *}\mid \xi_{kj}^{\nu *})q(\tau_{kj}^{\nu}\mid \tau_{kj}^{\nu *})}{p(\boldsymbol{\beta}_{kj}^{\nu};\tau_{kj}^{\nu})\pi({\tau_{kj}^{\nu}}^{2}\mid \xi_{kj}^{\nu})q({\tau_{kj}^{\nu *}}^{2}\mid \tau_{kj}^{\nu})}\right\}$$

We similarly update $(\tau_{kj}^{\eta}, \xi_{kj}^{\eta})$ in the case subclass regression.

6. Update the smoothness selection hyperparameter $\rho^{\nu}$ by

$$[\rho^{\nu}\mid others] \stackrel{d}{\sim} \mathsf{Beta}(a_{\rho}+\sum_{k,j}\mathbf{1}\{\xi_{kj}^{\nu}=1\}, b_{\rho}+\sum_{k,j}\mathbf{1}\{\xi_{kj}^{\nu}=0\}).$$

We similarly update $\rho^{\eta}$ in the case subclass weight regressions.

7. Update the scale parameter $\tau_{k0}$ in the hyperpriors for the intercept $\mu_{k0}^{*}$

$$[\tau_{k0}\mid others] \propto [\mu_{k0}^{*}\mid \tau_{k0}][\tau_{k0}\mid a_0, b_0] \stackrel{d}{\sim} \mathsf{Gamma}\left(a_0+\frac{1}{2}, b_0+\frac{(\mu_{k0}^{*})^2}{2}\right), k=1,\ldots,K-1.$$

8. Update the vector of subclass TPR for $j=1,...,J$ by

$$\begin{aligned}[\boldsymbol{\theta}^{(j)}\mid others] &\propto \prod_{\{i:\iota_{ij}=1\}}[\boldsymbol{M}_i\mid \boldsymbol{\theta}^{(j)}, Z_i, \boldsymbol{\iota}_i][\boldsymbol{\theta}^{(j)}]\\ &\propto \prod_{k=1}^{K}\left\{\theta_k^{(j)}\right\}^{m_{k1}^{(j)}}\left\{1-\theta_k^{(j)}\right\}^{m_{k0}^{(j)}}\cdot[\boldsymbol{\theta}^{(j)}],\end{aligned}$$

where $m_{kc}^{(j)} = \#\{i: Y_i=1, Z_i=k, \iota_{ij}=1, M_{ij}=c\}$, $c=0,1$. If prior for TPRs are independent Beta distributions, then this is a product of Beta distributions.

9. Update subclass-specific FPRs $\psi_k^{(j)}$ for $j=1,...,J$, $k=1,...,K$ from

$$\begin{aligned}[\psi_k^{(j)}\mid others] &\propto \prod_{i:\iota_{ij}=0, Z_i=k}[M_{ij}\mid \boldsymbol{\psi}^{(j)}, Z_i, I_i]\cdot[\psi_k^{(j)}]\\ &\propto \left\{\psi_k^{(j)}\right\}^{s_{k1}^{(-j)}}\left\{1-\psi_k^{(j)}\right\}^{s_{k0}^{(-j)}}\cdot[\psi_k^{(j)}],\end{aligned}$$

where $s_{kc}^{(-j)} = \#\{i : Z_i = k, \iota_{ij} = 0, M_{ij} = c\}$, for $c = 0, 1$. If the prior on FPRs are $\mathsf{Beta}(a_\psi, b_\psi)$, then the above conditional distribution is $\mathsf{Beta}(a_\psi + s_{k1}^{(j)}, b_\psi + s_{k0}^{(j)})$.

10. Update the regression parameters in the CSCF regression for $\ell = 1, \ldots, L$ by

$$
\begin{aligned}
[\Gamma_\ell^\pi \mid others] &\propto [\Gamma_\ell^\pi \mid \kappa_\beta, \kappa_\gamma] \cdot \prod_{i:Y_i=1} [I_i = \ell \mid \boldsymbol{\Gamma}^\pi, \boldsymbol{X}_i] \\
&\propto \prod_{i:Y_i=1} \frac{\{\exp(\alpha_{i\ell}^\pi)\}^{\mathbf{1}\{I_i=\ell\}}}{1+\exp(\alpha_{i\ell}^\pi)} \cdot \mathcal{N}(\boldsymbol{b}^\pi, \mathbf{K}^\pi),
\end{aligned}
$$

where $\boldsymbol{b}^\pi = 0$ is the prior mean, $\mathbf{K}_0^\pi = \mathrm{diag}\{\underbrace{\kappa_\beta, \ldots, \kappa_\beta}_{\sum_{j=1}^{p_1} p_1 \cdot C_j^\pi}, \underbrace{\kappa_\gamma, \ldots, \kappa_\gamma}_{p-p_1}\}$ is the prior precision matrix, and $\alpha_{i\ell}^\pi = \mathcal{X}_i^\top \boldsymbol{\Gamma}_\ell^\pi - D_{i\ell}$ with $D_{i\ell} = \log \sum_{\ell' \neq \ell} \exp(\mathcal{X}_i^\top \boldsymbol{\Gamma}_{\ell'}^\pi)$ with

$$
\mathcal{X}_i^\top = [\boldsymbol{B}_1^\top(X_{i1}), \ldots, \boldsymbol{B}_{p_1}^\top(X_{ip_1}), \widetilde{X}_{i,p_1+1}, \ldots, \widetilde{X}_{ip}].
$$

We introduce augmented Pólya-Gamma variables during sampling:

$$
\begin{aligned}
&[\omega_{i\ell}^\pi \mid \Gamma_\ell^\pi, others] \overset{d}{\sim} \mathsf{PG}(1, \alpha_{i\ell}^\pi), \\
&[\boldsymbol{\Gamma}_\ell^\pi \mid \omega_{i\ell}^\pi, others] \overset{d}{\sim} \mathcal{N}(\boldsymbol{m}_\ell^\pi, (V_\ell^\pi)^{-1}), \\
&V_\ell^\pi = \left\{\boldsymbol{\mathcal{X}}^\top \Omega_\ell^\pi \boldsymbol{\mathcal{X}} + \mathbf{K}^\pi\right\}^{-1}, \\
&\boldsymbol{m}_\ell^\pi = V_\ell^\pi \left\{\boldsymbol{\mathcal{X}}^\top (\boldsymbol{s}_\ell + \Omega_\ell^\pi D_\ell) + \mathbf{K}^\pi \boldsymbol{b}^\pi\right\},
\end{aligned}
$$

where $\boldsymbol{s}_\ell = \{s_{i\ell}\}_{i:Y_i=1}$, $s_{i\ell} = \mathbf{1}\{I_i = \ell\} - 1/2$, and $\Omega_\ell^\pi = \mathrm{diag}\{\{\omega_{i\ell}^\pi\}_{i:Y_i=1}\}$; We sample the PG variables in parallel across cases.

In simulations and data analysis, we ran three MCMC chains each with a burn-in period of 10,000 iterations followed by 10,000 iterations stored for posterior inference. We look for potential non-convergence in terms of Gelman-Rubin statistic (Brooks and Gelman, 1998) that compares between-chain and within-chain variances for each model parameter where a large difference ($R_c >$ 1.1) indicates non-convergence; We also used Geweke's diagnostic that compare the observed mean for each unknown variable using the first 10% and the last 50% of the stored samples where a large $Z$-score indicates non-convergence ($|Z| > 2$). In our simulations and data analyses, we observed fast convergence (many satisfied convergence criteria within $2,000$ iterations) that led to well recovered regression curves, TPRs and FPRs.

#### A4. Additional Information about Simulations of Main Paper

*Simulation I.* we let $\pi_\ell(\cdot)$, $\nu_k(\cdot)$ and $\eta_k(\cdot)$ depend on the two covariates $\boldsymbol{X} = \boldsymbol{W} = (S, T)$, $S$ and enrollment date ($T$), so that regression adjustments are necessary (see Remark 2 in the Main Paper). We simulate BrS measurements on $J = 9$ pathogens and assume the number of potential single-pathogen causes $L = J = 9$. To specify CSCF functions that satisfy the constraint $\sum_{\ell=1}^{L} \pi_\ell(\boldsymbol{x}) = 1$, we use stick-breaking parameterization with $L = 9$ segments. In particular, we let $\mathsf{logit}\ \{g_1(s,t)\} = \beta_1 \mathbf{1}(s = 1) + \sin(8\pi(t-0.5)/7)$, $\mathsf{logit}\ \{g_2(s,t)\} = \beta_2 \mathbf{1}(s = 1) + 4\exp(3t)/(1+\exp(3t)) - 0.5$, $\mathsf{logit}(g_\ell) = \beta_8 \mathbf{1}(s = 1)$ for $\ell > 2$; Let the CSCF functions $\pi_\ell(s,t) = g_\ell(s,t)\prod_{j<\ell}\{1 - g_j(s,t)\}, \ell = 1, \ldots, L(=9)$, where $\beta_\ell = 0.1, \ell = 1, \ldots, 8$. The true control distribution depend on covariates with $K = 2$ subclass weight functions: $\nu_1(s,t) = \mathsf{logit}^{-1}\{\gamma_1^\nu \mathbf{1}(s=1) + 4\exp(3t)/(1+\exp(3t)) - 0.5\}$ and $\nu_2(s,t) = 1 - \nu_1(s,t)$. We specify $\eta_k(s,t) = \nu_k(s,-t), k = 1, 2$, highlighting the need for using different subclass weights among cases and controls. We set the true TPRs $\theta_k^{(j)} = 0.95$ and the FPRs $\psi_1^{(j)} = 0.5$ and $\psi_2^{(j)} = 0.05$.

In performing regression analyses of the simulated data, we set $\phi_\ell(\boldsymbol{X})$ to be an additive model of a $\mathbf{1}\{S = 2\}$ indicator and a B-spline expansion with 7 degrees of freedom (d.f.) for standardized enrollment date $t$. We use $K = 7 (> K_0)$ and specify the regression formula for subclass weights $\nu_k(\cdot)$ and $\eta_k(\cdot)$ by additive models of the $\mathbf{1}\{S = 2\}$ indicator and a B-spline expansion with 5 d.f. for standardized enrollment date.

*Simulation II.* We consider $L = J = 3, 6, 9$ causes, under single-pathogen-cause assumption, BrS measurements made on $N_d$ cases and $N_u$ controls for each level of $X$ where $N_d = N_u = 250$ or

500. The functions $\phi_\ell(X) = \beta_{0\ell} + \beta_{1\ell}\mathbf{1}\{X=2\}$ take two sets of values to reflect how variable the CSCFs are across the two $X$ levels: i) $\boldsymbol{\beta}_0^{\mathsf{i}} = (0,0,0,0,0,0)$ and $\boldsymbol{\beta}_1^{\mathsf{i}} = (-1.5,0,-1.5,-1.5,0,-1.5)$ where causes have uniform CSCFs when $X=1$ and causes $\mathsf{B}$ and $\mathsf{E}$ dominate when $X=2$, or ii) $\boldsymbol{\beta}_0^{\mathsf{ii}} = (1,0,1,1,0,1)$ and $\boldsymbol{\beta}_1^{\mathsf{ii}} = (-1.5,1,-1.5,-1.5,1,-1.5)$ to mimic the scenario where pathogens $\mathsf{B}$ and $\mathsf{E}$ have lower CSCFs when $X=1$ and occupy more fractions when $X=2$. We further let the measurement error parameters take distinct values of the TPRs $\theta_k^{(j)} = 0.95$ or 0.8 and the FPRs $(\psi_1^{(j)}, \psi_2^{(j)}) \in \{(0.5,0.05),(0.5,0.15)\}$, for $j=1,\ldots,J$. Finally, we set the truth $\nu_k(W) = \eta_k(W) = \mathsf{logit}^{-1}(\gamma_{k0} + \gamma_{k1}\mathbf{1}\{W=2\})$ where $(\gamma_{10},\gamma_{11}) = (-0.5,1.5)$ and $(\gamma_{20},\gamma_{21}) = (1,-1.5)$. The Main Paper only presents results under $J=6$ for simplicity; The findings for $J=3,9$ are largely the same.

*Simulation* $\mathsf{II}$*: a randomly chosen replication.* Here we illustrate the inferences about the stratum-specific and overall CSCFs that are available to an analyst by considering a two-level covariate $X=W$ with $J=6$ measurements. Under the single-pathogen cause assumption, we can estimate $12 = (2\times 6)$ CSCFs, six per level of $X$ as well as six overall CSCFs. For example, based on a single data set simulated under the scenario $\{L=6,\ N_d=500,\ K=2,\ \theta_k^{(j)}=0.8,\ (\psi_1^{(j)},\psi_2^{(j)}) = (0.5,0.05),\ (\boldsymbol{\beta}_0^{\mathsf{ii}},\boldsymbol{\beta}_1^{\mathsf{ii}})\}$, Figure S2 in the Supplementary Materials shows the posterior distribution of the stratum-specific etiology fractions $\pi_\ell(X=s)$ for $(s=1,2)$ by row and $L(=J)$ causes $(\ell=1,\ldots,6)$ by column with the true values indicated by the blue vertical dashed lines; The bottom row shows the posterior distribution of $\pi_\ell^* = \sum_s w_s\pi_\ell(X=s)$ for $L$ causes with empirical weights $w_s = N_d^{-1}\sum_{i:Y_i=1}\mathbf{1}\{X_i=s\}$, $s=1,2$. The true stratum-specific and overall CSCFs are covered by their respective 95% CrIs.

#### A5. Additional Simulation Results

##### A5.1 *Estimating* $\pi_\ell(X)$

We use simulation studies to show the frequentist performance of the npLCM regression model in recovering stratum-specific CSCFs; The results below are based on a single discrete covariate that influence the CSCFs but not the subclass weights in the cases or controls.

In this simulation study, we simulate 500 cases and 500 controls for each of 7 sites. Every subject is measured on 6 pathogens A to F; The causes of disease are single-pathogen causes A-F. First, we let the CSCFs vary by site which are shown in Table S1. Second, we simulate the data using $K = 1$ subclass.

Table S1: True CSCFs for seven sites (boldfaced numbers indicate the highest CSCFs within each stratum).

| site\cause | A | B | C | D | E | F |
|---|---|---|---|---|---|---|
| 1 | **0.5** | 0.2 | 0.15 | 0.05 | 0.05 | 0.05 |
| 2 | 0.2 | **0.5** | 0.15 | 0.05 | 0.05 | 0.05 |
| 3 | 0.2 | 0.15 | **0.5** | 0.05 | 0.05 | 0.05 |
| 4 | 0.2 | 0.15 | 0.05 | **0.5** | 0.05 | 0.05 |
| 5 | 0.2 | 0.15 | 0.05 | 0.05 | **0.5** | 0.05 |
| 6 | 0.2 | 0.15 | 0.05 | 0.05 | 0.05 | **0.5** |
| 7 | 0.05 | 0.2 | 0.15 | **0.5** | 0.05 | 0.05 |

We simulate data under two TPR scenarios (I) strong signal with $\theta_1^{(j)} = 0.99$ and $\psi_1^{(1)} = 0.01$ where data are expected to provide strong information about the CSCFs, and (II) weak signal with $\theta_1^{(j)} = 0.55$ and $\psi_1^{(1)} = 0.45$ where it is easy to confuse true and false positive results and the data do not provide strong information about the CSCFs. In both scenario (I) and (II) , we used a Beta(6,2) distribution as a prior for the TPRs of the BrS measurements. We set the true TPRs and FPRs to be the same across sites and pathogens. In fitting the regression models, we use the CSCF regression formulation by specifying $L - 1$ sets of regression parameters with site dummy variables as the predictors in $\phi_\ell(\cdot)$. Since our goal is to infer $S = 7$ sets of CSCFs, we can also specify $S = 7$ sets of symmetric Dirichlet priors with hyperparameter $\alpha$ ($\mathsf{Dir}(\alpha)$); We use $\alpha = 1$ here. The package `baker` (`https://github.com/zhenkewu/baker`) provides an option to use Dirichlet priors when the CSCFs depend on discrete covariates only.

A5.1.1 *Scenario I: Strong Signal* Over $R = 100$ replications, the top half of Table S3 summarizes the coverage rates of the 95% credible intervals (CrIs) for the CSCFs across all the sites. We observed excellent recovery of the true values across all causes and sites with the 95% CrIs covered the true values between 90% to 100% of the time. Panel I of Table S3 also shows for site 1 the posterior mean CSCFs, posterior standard deviations (sd's) of the CSCFs, and posterior mean squared errors (PMSEs, estimated by $B^{-1}\sum_{b=1}^{B}\sum_{i:Y_i=1}\{\pi_\ell(X_i = s; \boldsymbol{\gamma}^{\pi,(b)}) - \pi_\ell^0(X_i = s)\}$ with $B$ retained posterior samples $\{\boldsymbol{\gamma}^{\pi,(b)}\}$) averaged over $R$ replications. The posterior means provide excellent estimation of the CSCFs with small average PMSEs.

A5.1.2 *Scenario II: Weak Signal* Using data simulated under less discrepant TPRs and FPRs than those in Scenario $\mathsf{I}$, the 95% CrIs cover the truths well for most site-cause pairs, but undercover the truths for causes with the highest CSCF in each site (see Table S2). This is expected because when the signal from the data is weak, the model relies more heavily on the uniform prior distribution for the CSCFs (symmetric Dirichlet prior with hyper-parameter 1).

Table S2: Number of times (out of 100 replications) that the true value is covered by the 95% CrIs (Scenario II, $\mathsf{Beta}(6,2)$ prior for the TPRs). Boldfaced numbers indicate the highest CSCFs (0.5) within each stratum.

| site\cause | A | B | C | D | E | F |
|---|---|---|---|---|---|---|
| 1 | **73** | 100 | 100 | 99 | 100 | 100 |
| 2 | 100 | **79** | 100 | 100 | 100 | 99 |
| 3 | 100 | 100 | **83** | 98 | 100 | 100 |
| 4 | 100 | 100 | 100 | **73** | 100 | 99 |
| 5 | 99 | 100 | 100 | 100 | **85** | 100 |
| 6 | 100 | 100 | 100 | 99 | 100 | **88** |
| 7 | 100 | 100 | 100 | **81** | 100 | 99 |

*More Informative TPR Priors ($II^*$).* We further investigate the model performance when we change the TPR prior distributions from the Beta(6,2) to a Beta distribution that has 95% of its mass between 0.525 and 0.575 and is around the true TPRs (Beta(835.95, 683.79), `beta_parms_from_quantiles(c(0.525,0.575))` using `baker`). Panel $II^*$ of Table S3 shows dramatic improvements in the coverage rates. These results suggest that changing the prior distributions of the TPRs so that it is more tightly concentrated around plausible values can improve inferences of the stratum-specific CSCFs in the presence of high levels of noises. Relative to Scenario I, the average PMSEs are larger across sites and pathogens reflecting the weaker signal in this setting.

In summary, in the simulation study where the CSCFs are influenced by a discrete covariate, the regression model recovers the true values well under high signals (high sensitivities and low FPRs). Under lower sensitivities and higher FPRs, the noisier simulated data are less informative about the CSCFs which are then more influenced by the prior distributions of the TPRs and CSCFs. In practice, we recommend eliciting quality informative TPR priors from domain scientists as in the PERCH study and perform sensitivity analyses to understand the robustness of the results with respect to the prior distributions.

##### A5.2 *Valid inference of $\pi^*_\ell$ omitting covariates*

Under the assumption in Remark 2 in the Main Paper, the case subclass weights $\boldsymbol{\eta}_k(\boldsymbol{W}) = \eta_k$, $k = 1, \ldots, K$, we conduct a simulation study to show that an npLCM analysis omitting covariates is able to provide valid inference about the overall CSCFs ($\boldsymbol{\pi}^*_\ell$). The simulation settings are exactly the same as in Simulation II, Section 5 in the Main Paper, except that we set $\gamma_{20} = \gamma_{21} = 0$ to satisfy the assumption. Figure S4(a) in the Supplementary Materials shows the percent relative biases are similarly negligible in all the 16 scenarios with 6 disease classes; Figure S4(b) in the Supplementary Materials shows excellent empirical coverage rates of the 95% CrIs for $\{\pi^*_\ell\}$.

Table S3: Scenario I and II*: coverage rates of the 95% CrIs; For Site 1, the posterior means, standard deviations (s.d.'s) and PMSE of the stratum-specific CSCFs averaged over $R = 100$ replications are also shown. Boldfaced numbers indicate the highest CSCFs (0.5) within each stratum.

| | | site \cause | A | B | C | D | E | F |
|---|---|---|---|---|---|---|---|---|
| I | coverage | 1 | **99** | 93 | 97 | 94 | 96 | 90 |
| | | 2 | 97 | **90** | 96 | 97 | 95 | 94 |
| | | 3 | 100 | 95 | **98** | 98 | 95 | 96 |
| | | 4 | 93 | 94 | 96 | **95** | 92 | 99 |
| | | 5 | 96 | 94 | 96 | 97 | **95** | 98 |
| | | 6 | 96 | 97 | 98 | 99 | 95 | **96** |
| | | 7 | 96 | 97 | 91 | **100** | 95 | 96 |
| | posterior summary | truth (Site 1) | 0.5 | 0.2 | 0.15 | 0.05 | 0.05 | 0.05 |
| | | average of post. mean | 0.495 | 0.197 | 0.152 | 0.053 | 0.053 | 0.051 |
| | | average of post. s.d. | 0.023 | 0.018 | 0.016 | 0.01 | 0.01 | 0.01 |
| | | average PMSE | 0.0010 | 0.0007 | 0.0005 | 0.0002 | 0.0002 | 0.0002 |
| II* | coverage | 1 | **98** | 89 | 98 | 99 | 100 | 100 |
| | | 2 | 97 | **95** | 96 | 100 | 100 | 99 |
| | | 3 | 93 | 98 | **91** | 99 | 99 | 100 |
| | | 4 | 95 | 98 | 100 | **95** | 99 | 100 |
| | | 5 | 94 | 94 | 99 | 99 | **91** | 100 |
| | | 6 | 95 | 97 | 100 | 99 | 99 | **90** |
| | | 7 | 100 | 95 | 94 | **96** | 100 | 99 |
| | posterior summary | truth (Site 1) | 0.5 | 0.2 | 0.15 | 0.05 | 0.05 | 0.05 |
| | | average post. mean | 0.417 | 0.163 | 0.138 | 0.091 | 0.086 | 0.106 |
| | | average post. s.d. | 0.27 | 0.174 | 0.162 | 0.135 | 0.13 | 0.141 |
| | | average PMSE | 0.131 | 0.067 | 0.056 | 0.034 | 0.031 | 0.042 |

#### A6. Results from PERCH Data: Individual-level Probability Assignment

The regression model accounts for stratification of CSCFs by the observed covariates. Consequently, despite identical diagnostic test results, the posterior algorithm automatically assigns covariate-dependent cause-specific probabilities. Figure S5 shows the individual cause-specific probabilities for cases with all negative NPPCR results (the most frequent pattern among cases). For two cases with one older the other younger than one, the older case has a lower posterior probability of her disease caused by RSV and higher probability of being caused by NoS. Indeed, contrasting older and younger children while holding the enrollment date, HIV, severity constant, the estimated difference in the log odds (i.e., log odds ratio) of a child being caused by RSV versus

NoS is negative: $-1.82$ (95% $\mathrm{CrI}: -2.99, -0.77$).

#### A7. Supplemental Figures

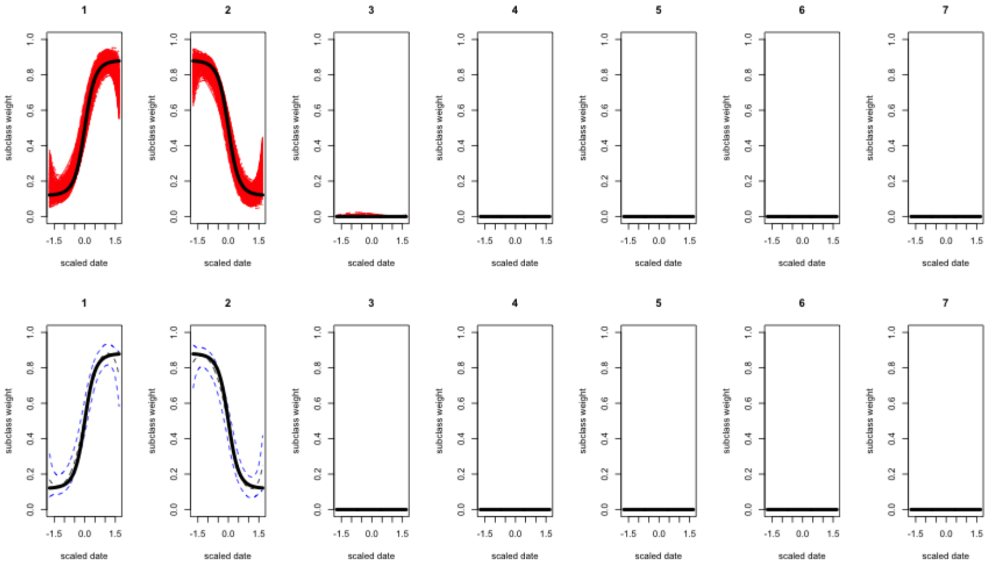


(a) case

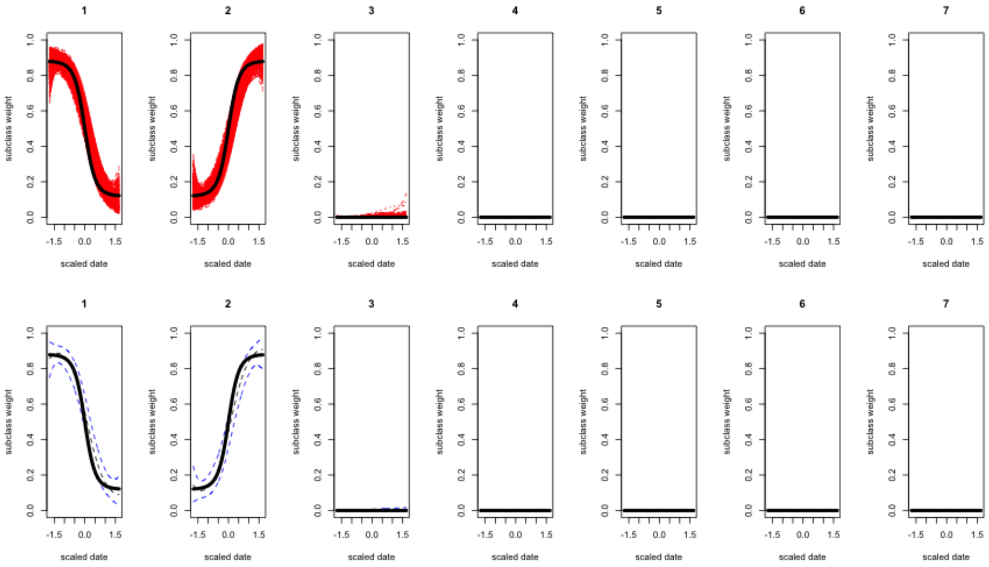


(b) control

Fig. S1: By propagating the prior that encourages few subclasses, the algorithm correctly infers two subclasses from the simulated data in Simulation I, Section 5 in the Main Paper. Estimated case (top) and control (bottom) subclass weight curves over one continuous covariate $\widehat{\nu}_k(t)$ (central blue dashed lines enclosed by the 95% credible regions; the red curves are posterior samples) compared against the simulation truths ($\nu_k^0(t)$, black solid lines). The number of subclasses is bounded by seven during model fitting.

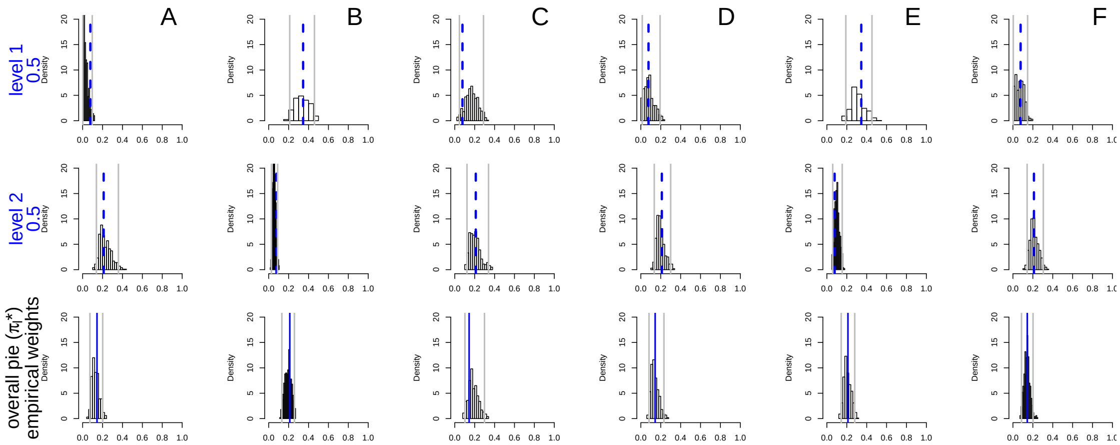


Fig. S2: Posterior distributions of the stratum-specific (Row 1 and 2) and the overall (Bottom Row) CSCFs based on a simulation with a two-level discrete covariate and $L = J = 6$ causes. The vertical gray lines indicate the 2.5% and 97.5% posterior quantiles, respectively; The truths are indicated by vertical blue dashed lines. *Row 1-2*) CSCFs by stratum (level = 1,2) and cause (A-F); *Bottom*) $\pi^*_\ell$: overall population etiologic fraction for cause A-F (empirical average of the two CSCFs above).

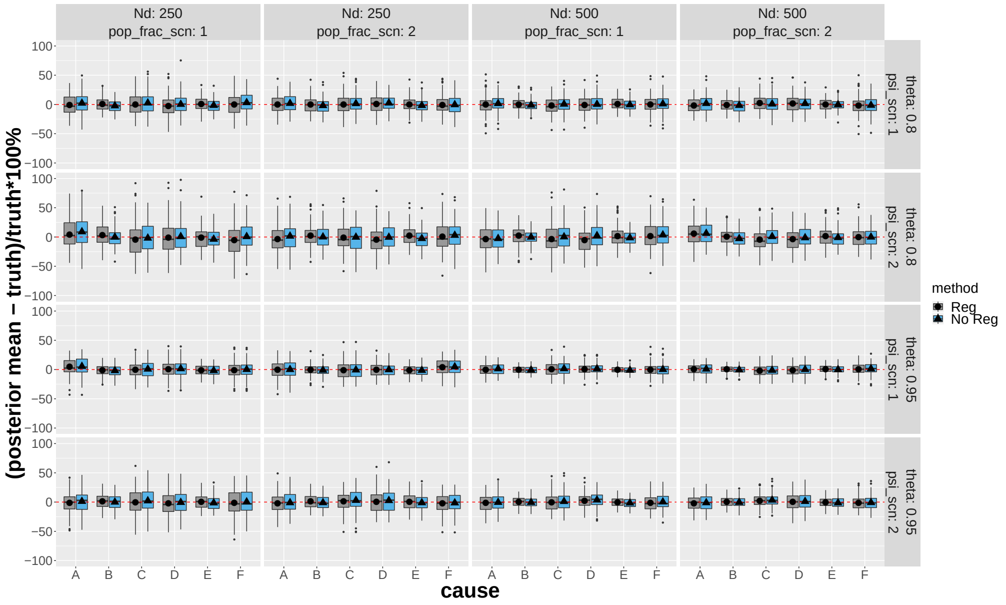


(a)

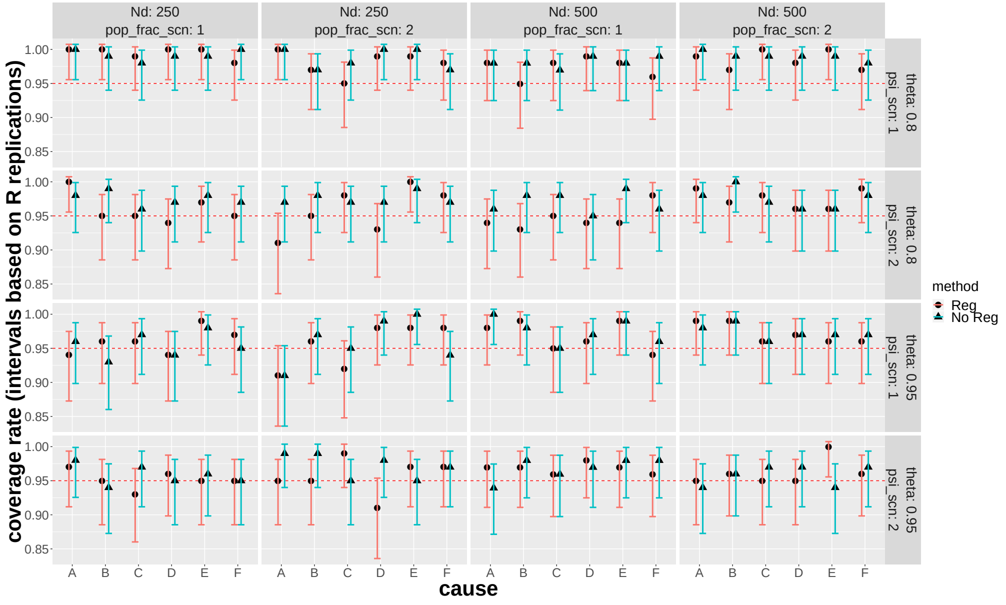


(b)

Fig. S3: NPLCM analyses with or without regression perform similarly in terms of percent relative bias (top) and empirical coverage rates (bottom) over $R = 100$ replications in simulations where the case and control subclass weights *do not* vary by covariates. Each panel corresponds to one of 16 combinations of true parameter values and sample sizes. See Figure 4 in the Main Paper for detailed descriptions of the figure.

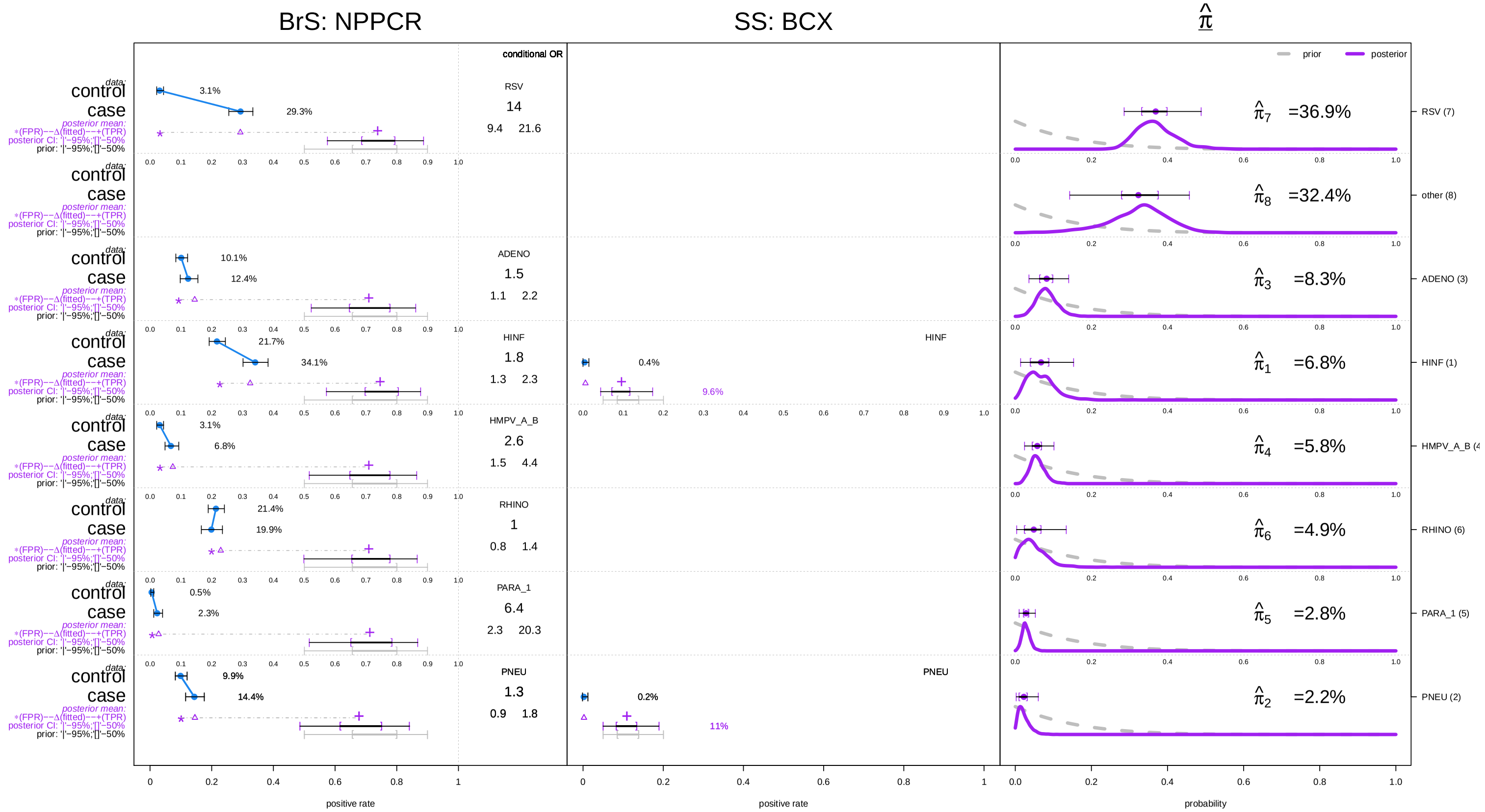


Fig. S4: Panel plot with NPPCR (Bronze-Standard, BrS), BCX (Silver-Standard, SS) and Etiology Pies obtained from an npLCM analysis omitting covariates ($K = 5$). For each of the 7 pathogens, a summary of the NPPCR and BCX data analyzed in Section 6 of Main Paper is shown in the left two columns, along with some of the intermediate model results; and the prior and posterior distributions for the CSCFs on the right (rows ordered by posterior means). *Left*) The observed BrS rates (with 95% confidence intervals, CI) for cases and controls are shown on the far left with solid dots. The conditional odds ratio (COR) contrasting the case and control rates given the other pathogens is listed with 95% CI in the box to the right of the BrS data summary. Below the case and control observed rates is a horizontal line with a triangle. From left to right, the line starts at the estimated false positive rate (FPR, $\widehat{\psi}_j^{\mathsf{BrS}}$) and ends at the estimated true positive rate (TPR, $\widehat{\theta}_j^{\mathsf{BrS}}$), both obtained from the model. Below the TPR are 95% and 50% intervals summarizing its posterior (top) and prior (bottom) distributions for that pathogen. These intervals show how the prior assumption influences the TPR estimate as expected given the identifiability constraints. The triangle on the line is the model estimate of the case rate to compare to the observed value above it. *Middle*) The SS data are shown in a similar fashion to the right of the BrS data. By definition, the FPR is 0.0 for SS measures and there is no control data. The observed rate for the cases is shown with its 95% CI. The estimated SS TPR ($\widehat{\theta}_j^{\mathsf{SS}}$) with prior and posterior distributions is shown as for the BrS data. *Right*) The marginal posterior and prior distributions of the etiologic fraction for each pathogen. We appropriately normalized each density to match the height of the prior and posterior curves. The posterior mean with 50% and 95% CrIs are shown above the density.

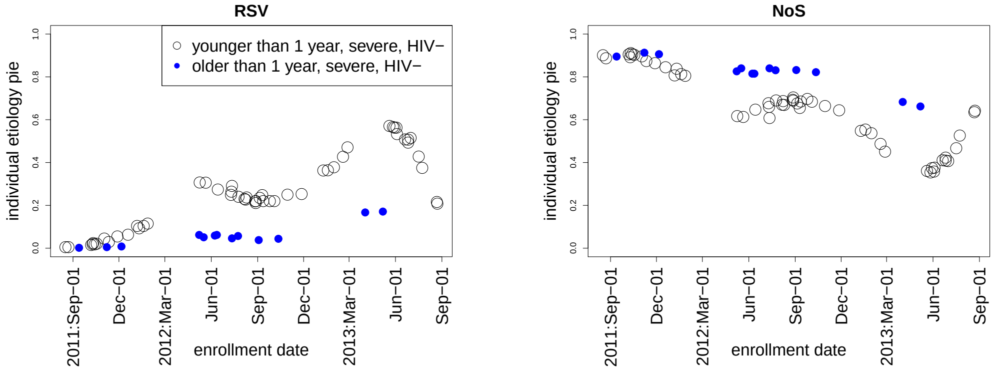


(a) Cause: RSV

(b) Cause: NoS

Fig. S5: Individual-level probabilities of cause of pneumonia estimates for RSV (left) and NoS (right) differ by age and season among HIV negative and severe pneumonia cases for whom the seven pathogens were *all tested negative* in the nasopharyngeal specimens. The prediction also outputs probabilities for other causes; only RSV and NoS are shown here.

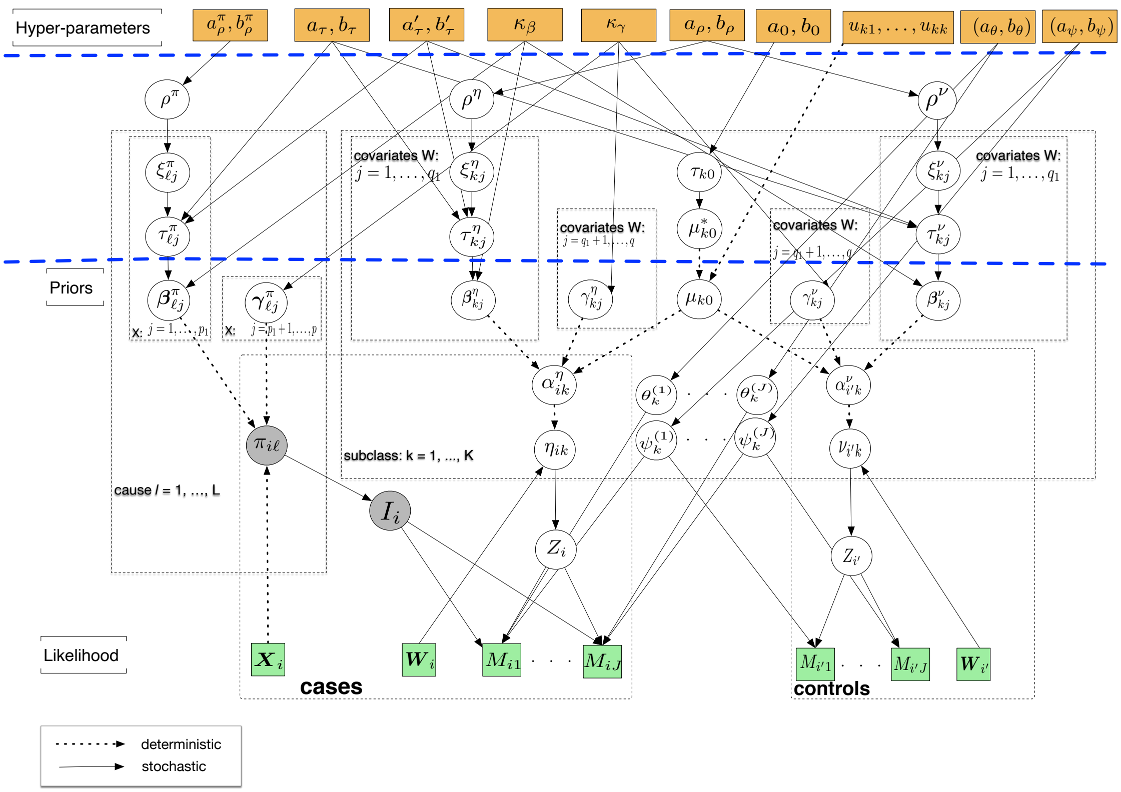


Fig. S6: The directed acyclic graph (DAG) representing the structure of the model likelihood and prior. The quantities in squares are either data or hyperparameters; the unknown quantities are shown in the circles. The arrows connecting variables indicate that the parent parameterizes the distribution of the child node (solid lines) or completely determines the value of the child node (dotted arrows). The rectangular "plates" where the variables are enclosed indicate that a similar graphical structure is repeated over the index; The index in a plate indicate subjects, causes, covariates or subclasses.
